# A BRCT Domain-Containing Protein Induced in Early Phagocytosis Plays a Crucial Role in Mucorales Pathogenesis

**DOI:** 10.1101/2025.05.13.652844

**Authors:** Ghizlane Tahiri, Carlos Lax, Ulrike Binder, Jakob Scheler, Eusebio Navarro, Francisco E. Nicolás, Victoriano Garre

## Abstract

Mucormycosis, caused by Mucoralean fungi, is among the most lethal fungal diseases, and a deeper understanding of its pathogenesis is urgently needed. Transcriptomic profiling of virulent (WT) and an RNAi-deficient strain (*r3b2*Δ) of *M. lusitanicus* strains during phagocytosis uncovered thousands of differentially expressed genes (DEGs), highlighting early metabolic activation as a key survival strategy inside the phagosome. Enriched pathways included amino acid transport, nucleotide metabolism, and translation, reflecting an adaptive fungal response to nutrient deprivation and host immune stress. Integrative analyses of mRNA and sRNA profiles also revealed a critical role of the RNAi pathways in modulating gene expression during infection.Building on these observations, we identified four chromatin- and transcription-related candidate virulence genes—*brca1*, *box*, *hist1*, and *hda10*—which were strongly upregulated during phagocytosis and regulated by RNAi. Functional validation through gene deletion in *M. lusitanicus* and disruption in *R. microsporus* revealed that while loss of these genes in *M. lusitanicus* did not significantly affect virulence, *R. microsporus* mutants for *brca1*, *hist1*, and *hda10* showed attenuated virulence in a murine model. Our findings suggest that although *M. lusitanicus* remains a valuable tool for genetic manipulation, species-specific differences must be considered when studying virulence. The study also underscores the importance of using multiple Mucorales models to uncover conserved and divergent strategies employed by pathogenic fungi. These insights contribute to a broader understanding of fungal adaptation, immune evasion, and the identification of novel targets for antifungal intervention.

## INTRODUCTION

Macrophages are essential phagocytes in the innate immune system, playing a crucial role in defending against invasive fungal infections. These immune cells are rapidly recruited to infection sites, where they recognize pathogen-associated molecular patterns (PAMPs) and phagocytose fungal spores (1). In lung infections of healthy mice and rats caused by fungal pathogens, alveolar macrophages effectively engulf spores and inhibit their germination through nitric oxide production and nutrient deprivation—mainly by restricting iron availability—ultimately leading to pathogen clearance. Once recognized by pattern recognition receptors (PRRs), fungal spores are internalized, forming phagosomes that subsequently fuse with lysosomes to create an acidic phagolysosome (2).

However, some fungal pathogens have evolved mechanisms to evade or manipulate this immune response, ensuring their survival inside phagocytes. Among them, *Mucorales* fungi—such as *Mucor* and *Rhizopus* species—are opportunistic pathogens responsible for mucormycosis, a severe and often fatal infection that primarily affects immunocompromised individuals (3). These fungi employ diverse strategies to resist or exploit phagocytosis, allowing them to persist within host cells and spread.

In vitro studies indicate that while macrophages can recognize and phagocytose Mucorales spores, certain species are able to germinate intracellularly, thereby surviving the host defense mechanisms (4). For instance, *Rhizopus* species can arrest phagosome maturation, preventing macrophage-mediated killing (5). Additionally, *Rhizopus microsporus* harbors the bacterial endosymbiont *Ralstonia pickettii*, which secretes factors that impair the ability of the host to detect and eliminate the fungal pathogen (6). Similarly, *Mucor lusitanicus* spores resist intracellular elimination by blocking phagosome maturation through a calcineurin-dependent mechanism, which promotes their survival and germination within macrophages (7). This ability to grow inside host cells is partially regulated by the transcription factors Atf1 and Atf2 (8). Furthermore, in *M. lusitanicus*, spore size influences virulence, as only larger spores can germinate inside macrophages. Interestingly, during zebrafish infections, *M. lusitanicus* induces apoptosis in macrophages but not in neutrophils (9).

During the interaction between *Mucorales* and their hosts, extensive genetic regulatory changes occur, including modulation of gene expression by RNA interference (RNAi). In *M. lusitanicus*, two major RNAi pathways have been described: the canonical RNAi pathway and the non-canonical RNAi Pathway (NCRIP) (10). The canonical pathway depends on Dicer and Argonaute proteins to process double-stranded RNA (dsRNA) into small interfering RNAs (siRNAs), which regulate gene expression and maintain genome stability (11).

In contrast, NCRIP functions independently of Dicer and Argonaute. Instead, it relies on the RNase III-like enzyme R3B2, which specifically cleaves single-stranded RNAs (ssRNAs), along with RNA-dependent RNA polymerases (RdRPs) (8). This pathway plays a crucial role in controlling gene expression under stress conditions, including host-pathogen interactions. Studies have shown that NCRIP is involved in fungal virulence, as its disruption alters stress responses and reduces pathogenicity, suggesting its importance in infection and survival within the host (11).

Previous studies have characterized the NCRIP pathway in fungal biology and pathogenesis, though they primarily focused on mRNA profiling during later stages of phagocytosis, once fungal spores had germinated (12). Here, we have performed a deeper characterization by conducting a simultaneous degradome analysis, integrating sRNA and mRNA profiles to explore gene regulation during macrophage infection. This comprehensive approach revealed coordinated control by canonical and NCRIP pathways, with *M. lusitanicus* modulating gene degradation in response to phagocytosis—highlighting molecular responses critical for fungal fitness and host adaptation. Additionally, we examined conserved virulence factors in *M. lusitanicus* and *R. microsporus*, focusing on early phagocytic interactions. Furthermore, this study investigates the role of conserved virulence factors in *M. lusitanicus* and *R. microsporus* by analyzing early phagocytosis events using mutant strains and infection models. Through functional analysis, we identified the *histone 1* ortholog as a key contributor to fungal fitness and the *brca1* ortholog as a determinant of pathogenesis. These findings provide novel insights into the genetic basis of virulence attenuation in *R. microsporus*, shedding light on conserved mechanisms of immune evasion during the initial stages of host-pathogen interaction.

## RESULTS

### *M. lusitanicus* deploys a major transcriptional response in early phagocytosis

To determine the transcriptional profile of *M. lusitanicus* spores during early phagocytosis, a time-course experiment was conducted at different intervals of macrophage internalization (30 min, 60 min, 90 min, 2 h, 3 h, and 5 h). The goal was to identify the most suitable time point at which the more substantial number of *M. lusitanicus* spores are phagocytosed, ensuring sufficient fungal RNA concentration for analysis. Two *M. lusitanicus* strains were used to monitor phagocytosis: a wild-type strain (WT, MU636) and a knockout mutant strain in the *r3b2* gene (MU412). The internalization process was analyzed using an in vitro host-pathogen system, where both strains were confronted with the J774A.1 mouse macrophages cell line (ATCCT1B-G7) at a ratio 1.5:1, respectively. As non-interaction controls, fungal spores from both strains were grown under saprophytic conditions in L15 medium (Figure 1A). mRNAs and sRNAs were extracted at each time point, as well as from the non-interaction controls. Although fungal RNA was detected at all time points, the one-hour mark (60 min) was selected for subsequent experiments, as nearly all fungal spores had been phagocytosed by this time (Figure 1B,C, D).

**Fig. 1.**
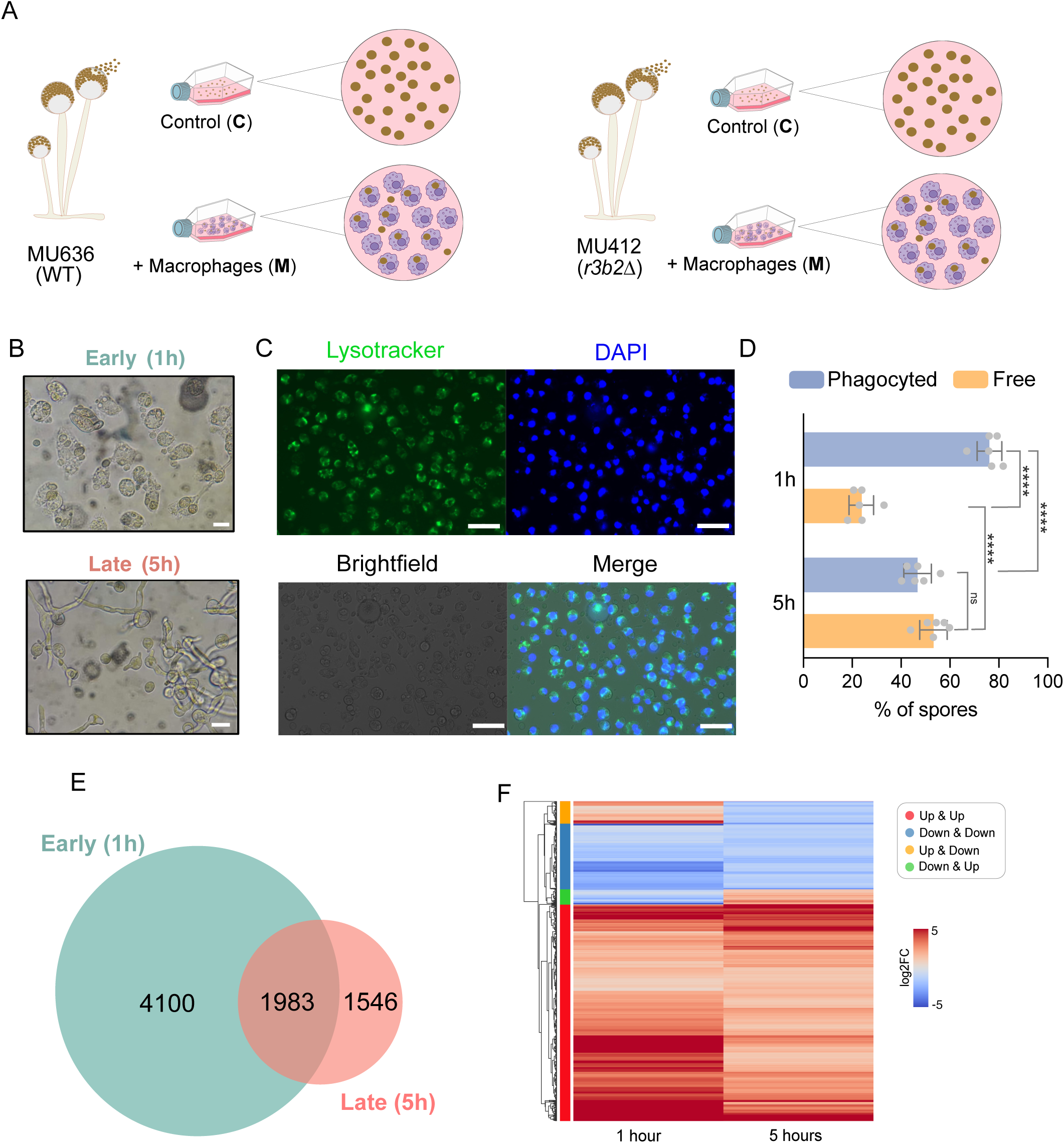
(**A)** Schematic representation of the RNA-seq analysis experimental design, including macrophage phagocytosis and saprophytic conditions. The assays included wild-type (WT) spores with and without macrophages (WT-M and WT-C), and *r3b2*Δ spores with and without macrophages (*r3b2*Δ-M and *r3b2*Δ-C). **(B)** Macrophages stained with LysoTracker Green DND-26 and Hoechst 3342 (blue) during early and late interactions with *Mucor* spores (scale bar = 15 µm). **(C)** Quantification of internalized versus free spores during early and late stages of phagocytosis (scale bar = 50 µm). Data are represented as mean ± SD. Statistical analysis was performed using Welch’s t-test. Significant differences were observed between phagocytosed and free spores at 1 h (p < 0.0001), and between phagocytosed spores at 1 h versus 5 h (p < 0.0001), as well as between free spores at 1 h and 5 h (p < 0.0001). No **significant** difference was found between free and phagocytosed spores at 5 h (p = 0.0711). **(D)** Comparative transcriptomic analysis of the wild-type *Mucor* strain during early and late phagocytosis. **(E)** Differentially expressed genes (DEGs) shared between early and late phagocytosis, classified by expression patterns: red indicates genes upregulated at both time points; blue, downregulated at both; orange, upregulated early and downregulated late; and green, downregulated early and upregulated late.

To investigate the transcriptional dynamics of *M. lusitanicus* during phagocytosis, we compared the DEGs of the WT strain in saprophytic conditions and after 1 hour (early phagocytosis) and 5 hours (late phagocytosis) of interaction. Interestingly, we identified 4100 genes that are exclusively differentially expressed during early phagocytosis, whereas only 1546 genes are uniquely expressed in the late phase (Figure 1E). These findings suggest that *M. lusitanicus* mounts a major transcriptional response during the initial stages of phagocytosis. The transcriptional profile comparison further revealed that 1983 genes are differentially expressed at both 1 hour and 5 hours post-phagocytosis (Figure 1F, Supplementary File 1). Among these, a subset of genes exhibits a chronological transcriptional response, with 1419 genes (Figure 1F, red) consistently upregulated and 473 genes (Figure 1E, blue) persistently downregulated at both time points. Additionally, some genes display a temporal activation specific to either the early or late phase of phagocytosis: 114 genes undergo transient upregulation at 1 hour followed by downregulation at 5 hours (Figure 1E, C3, orange), whereas 106 genes follow the opposite pattern, being downregulated at 1 hour and upregulated at 5 hours (Figure 1E, green).

### Transcriptional response of *M. lusitanicus* to early phagocytosis

To further dissect the transcriptional regulation during early phagocytosis, we expanded the analysis to include both the wild-type (WT) and RNAi-deficient *r3b2*Δ mutant strains. This allowed us to identify genes regulated by early phagocytosis, those controlled by RNAi, either through the NCRIP and/or the canonical RNAi pathway during this early stage, and genes that are simultaneously influenced by both RNAi and phagocytosis. For the WT strain, DEGs were identified by comparing macrophage interaction (WT-M) to saprophytic growth (WT-C) at 1 hour. In parallel, the same comparison was conducted for the *r3b2*Δ mutant (*r3b2*Δ-M vs *r3b2*Δ-C). These analyses revealed a total of 6083 differentially expressed genes (DEGs) in the WT strain (|log2FC ≥1|, FDR < 0.05) regulated by phagocytosis (Figure 2A) (Supplementary File 1). In parallel, the *r3b2*Δ strain showed a comparable number of DEGs—6167 genes—regulated during phagocytosis (*r3b2*Δ-M vs *r3b2*Δ-C) (Figure 2A) (Supplementary File 1). Notably, 5407 genes were commonly regulated by phagocytosis across both strains (Figure 2B). Interestingly, 760 genes were differentially expressed exclusively during the interaction of the attenuated *r3b2*Δ strain with macrophages, while 676 genes showed expression changes during the internalization of the virulent WT strain (Figure 2B). Additionally, 2778 commonly differentially expressed genes during phagocytosis of both strains were found to be upregulated. Together, these results suggest a strong interplay between the RNAi pathways in *M. lusitanicus* and the transcriptional regulation of phagocytosis response.

**Fig. 2.**
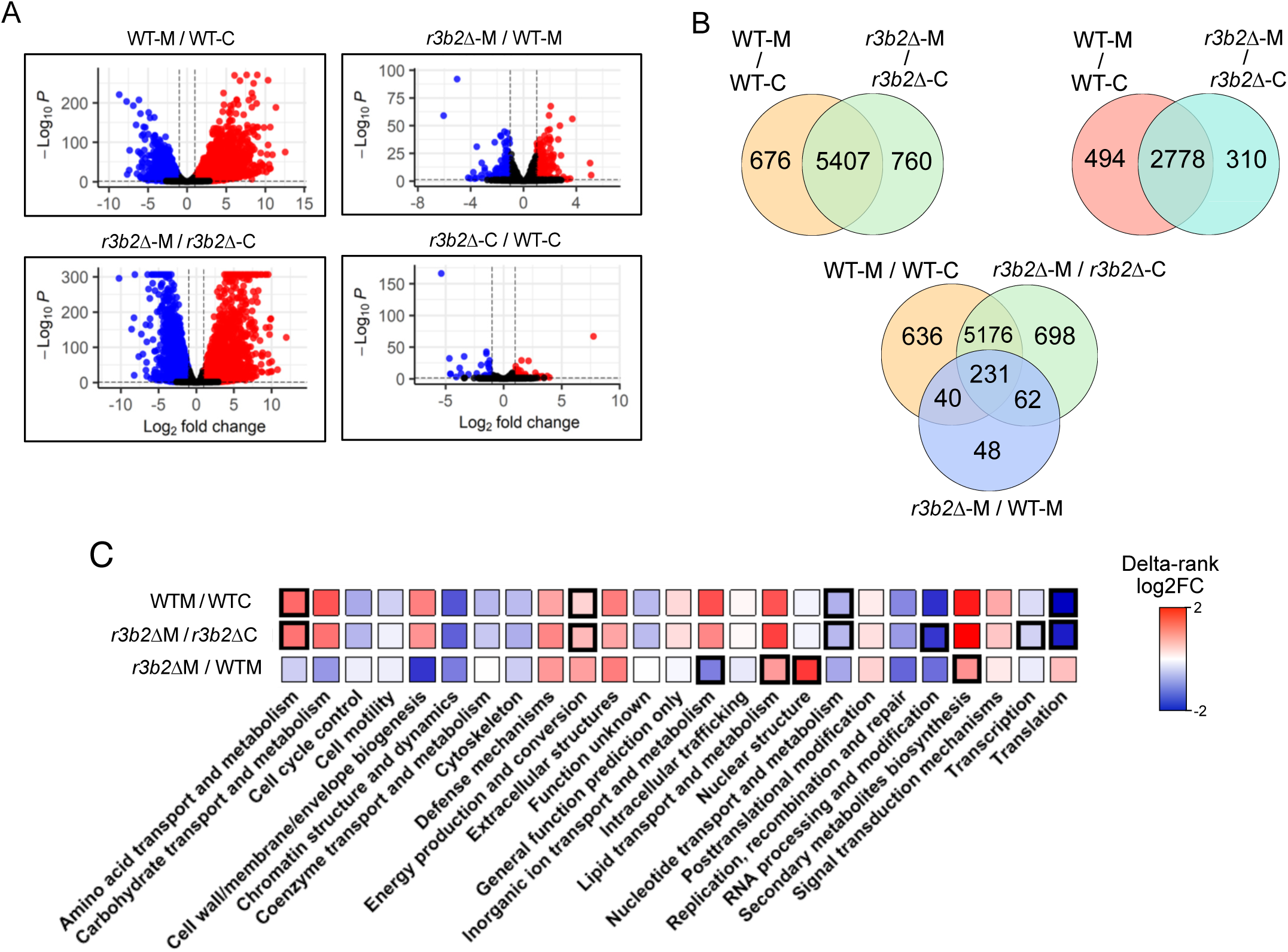
Early transcriptomic changes of the fungal spores during their phagocytosis by macrophages. **(A)** Volcano plots of the WT strain fungal spores with and without macrophages (WT-M/WT-C), the *r3b2Δ* versus the WT interacting with macrophages (*r3b2Δ*-M/ WT-M), the *r3b2Δ* with and without macrophages (*r3b2Δ*-M/ *r3b2Δ*-C), and the *r3b2Δ* versus the WT in saprophytic conditions (*r3b2Δ*-C/ WT-C). Red and blue dots indicate the upregulated and downregulated genes, respectively, while black dots show not differentially expressed genes. **(B)** Venn diagrams of the common DEGs of the fungal WT and mutant strains interacting with macrophages. The two set of the WT and the mutant fungal strains during their phagocytosis. The three set venn diagram reflects the common DEGs of the WT and the mutant strains interacting with macrophages and those resulted from the comparison of the mutant and the WT strains during their phagocytosis. **(C)** Enrichment analysis of the DEG of the WT during its interaction with macrophages versus its saprophytic growth, the enrichment analysis of those DE during the interaction of the mutant with macrophages versus its growth under non-stressful conditions, and the enrichment of the DEG when comparing the mutant versus the WT during their phagocytosis.

### The NCRIP pathway is active in the early phases of fungal development

Despite its implication in various biological processes (8) until now, however, it remained unclear whether this pathway is activated during the first stage of the fungal life cycle, particularly at the spore stage, and which genes it regulates during this phase. To address this, we compared the *r3b2*Δ mutant to the WT strain in a non-infection context (*r3b2*Δ-C vs WT-C) to identify genes controlled by the NCRIP in the early stages of the life cycle of the fungal pathogen (Figure 2A). Differential expression analysis revealed that, even at this early stage of development, the NCRIP pathway is active, regulating the expression of 132 genes after one hour of growth under non-stressful conditions (Figure 2A) (Supplementary File 1).

### Deciphering the gene network controlled by the virulence-related RNAi mechanism during early phagocytosis

Having demonstrated the activity of the NCRIP pathway during the initial stages of fungal development, specifically at the spore stage, which is the common morphology typically inhaled by hosts we wanted to asses whether the NCRIP pathway is active during the early phagocytosis of *M. lusitanicus* spores. To that end, we compared the transcriptional profiles of the *r3b2*Δ and WT strains during macrophage interaction. This analysis focused on DEGs shared between the WT-M vs WT-C and *r3b2*Δ-M vs WT-M comparisons, which yielded 381 DEGs (Figure 2A, Supplementary File 1), as well as those identified in the *r3b2*Δ-M vs *r3b2*Δ-C comparison. 5176 genes were commonly regulated by early phagocytosis regardless of the virulence of the phagocytosed strain, which could be involved in recognizing fungal external structures or molecules released by the fungus (Figure 2B). Additionally, 62 genes are regulated by both early phagocytosis of the WT strain and the NCRIP. These genes are differentially expressed during the interaction of the virulent WT strain with macrophages and are also regulated by the NCRIP, potentially contributing to invasion. Finally, 231 genes are co-regulated by both the NCRIP and phagocytosis, irrespective of strain virulence, indicating genes regulated by both processes.

To explore the functional categories of these gene sets, we performed a KOG analysis (Figure 2C). Results revealed that the NCRIP governs key metabolic processes, both under saprophytic conditions and during macrophage interaction—reinforcing its relevance in supporting virulence through metabolic adaptation. Even at the spore stage and under phagocytic stress, the NCRIP contributes to controlling core energy production mechanisms. Similarly, early phagocytosis itself regulates functions such as amino acid transport and metabolism, energy production, nucleotide metabolism, and translation. In the *r3b2Δ* mutant, comparison between saprophytic and phagocytosed conditions showed enrichment in transcriptional regulation and RNA processing. Biological processes co-regulated by both the NCRIP and early phagocytosis include inorganic ion and lipid transport, nuclear structure, and secondary metabolite biosynthesis, all of which may play key roles in establishing infection.

### Integrative degradome analysis: sRNA production and mRNA dynamics in the early phagocytosis of *M. lusitanicus*

While mRNA analysis enables the detection of genes regulated by the NCRIP—based on differential expression between the WT and *r3b2Δ* strains, it does not provide conclusive evidence of direct degradation by this mechanism or whether changes in gene expression are secondary to the degradation of upstream regulators. To address this, we conducted parallel sRNA sequencing on the same samples to identify direct targets of the NCRIP and genes regulated by other RNAi pathways. The analysis of sRNA data revealed that when the mutant strain interacts with macrophages, the number of differentially degraded genes (DDGs) reaches 2101 (*r3b2*Δ-M vs *r3b2*Δ-C, Supplementary File 2), compared to its saprophytic growth (Figure 3A). This is 3.5 times higher than the number observed in the WT strain during phagocytosis compared to its saprophytic growth (600 DDGs, WT-M vs WT-C, Supplementary File 2). The sRNAs produced in the mutant during its interaction with macrophages, relative to saprophytic growth, are not products of R3B2 activity, indicating that the canonical RNAi pathway is activated during early phagocytosis. Additionally, the higher number of DDGs in *r3b2*Δ-M vs *r3b2*Δ-C compared to WT-M vs WT-C suggests that the NCRIP may function as a negative regulator of the canonical RNAi pathway.

**Fig. 3.**
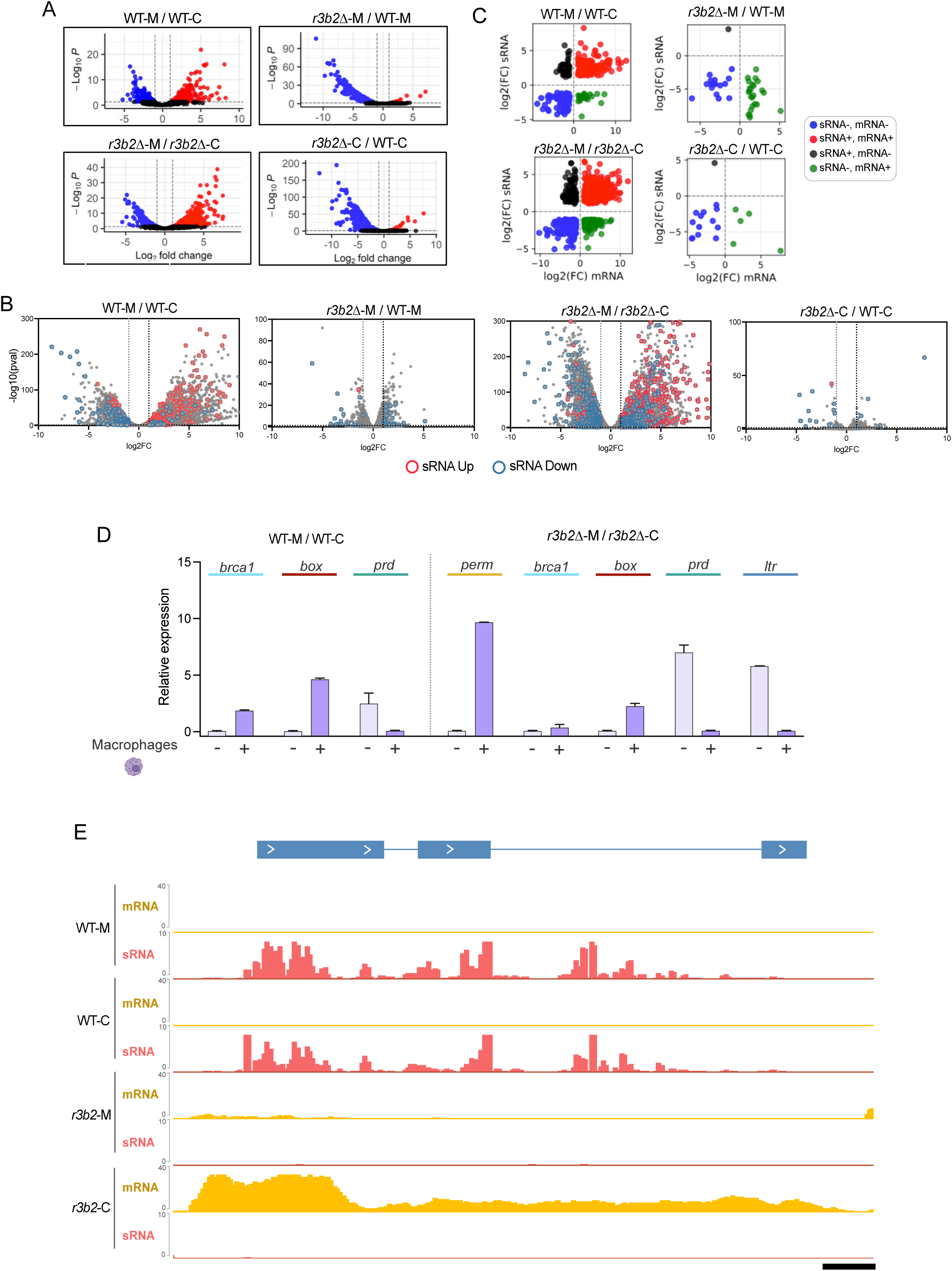
The degradome analysis based on the correspondence between the sRNA production and mRNA expression. **(A)** Volcano plots of the sRNA analysis of the WT confronted with macrophages versus its saprophytic growth., the analysis of the sRNA production comparing the mutant interacting with macrophages versus the WT interacting with macrophages, the analysis of the sRNA in the mutant with and without macrophages, and the study of the sRNA in the mutant versus the WT in non-stressful conditions. **(B)** Scatter plots showing the relationship between the expression of the sRNA and mRNA in the WT-M / WT-C, *r3b2Δ-*M / WT-M, *r3b2Δ-*M / *r3b2Δ-*C, and *r3b2Δ-*C / WT-C. Blue dots are genes with downregulation in their mRNA and sRNA, black dots are genes upregulated at the sRNA level and downregulation at the mRNA level, the red dots are genes with simultaneous upregulation in the sRNA and mRNA, and the green dots are genes with downregulated sRNA and upregulated mRNAs. **(C)** Analysis of differentially expressed genes (DEGs) from the mRNA dataset based on their corresponding sRNA levels across the following comparisons: WT-M / WT–C, *r3b2*Δ-M / WT-M, *r3b2*Δ-M / *r3b2*Δ–C, and *r3b2*Δ–C / WT–C. The number of upregulated and downregulated sRNAs was determined for each condition. **(D)** Validation of RNA-seq data by qRT-PCR. Gene expression was evaluated in WT spores interacting with macrophages versus saprophytic growth, and in the *r3b2*Δ mutant under the same conditions. Target genes included a permease, three transcription factors (*prd*, *box*, and *brca1*), and an LTR-transposable element. **(E)** Genomic coverage of sRNA and mRNA reads mapped to the LTR-transposon and adjacent genes in WT and *r3b2*Δ strains, under both saprophytic and macrophage-interacting conditions. Yellow and red plots represent sRNA and mRNA read coverage, respectively.

Additionally, the production of sRNAs resulting from the degradation of target mRNAs by the NCRIP was slightly higher during the saprophytic growth of the pathogen (742 DDGs in *r3b2*Δ-C vs WT-C, Supplementary File 2) compared to the interaction of fungal spores with macrophages (680 DEGs in *r3b2*Δ-M vs WT-M, Supplementary File 2) (Figure 3A). Notably, sRNA abundance was consistently higher in the WT strain compared to the mutant strain, both in the infection context (*r3b2*Δ-M vs WT-M) and under saprophytic conditions (*r3b2*Δ-C vs WT-C), suggesting a predominant role of NCRIP in regulating these genes.

Next, to determine the global correspondence between sRNA production and mRNA degradation plotted DEG genes at mRNA level, indicating which ones had a significant increase or decrease in sRNA accumulation (Figure 3B). Overall, we found a higher number of genes with significant changes in sRNA accumulation in the comparison in which a higher number of DEG were detected (Figure 3B). We also found that for all growth conditions, genes accumulating a higher number of sRNAs showed lower expression levels and the opposite happened with the set of genes that accumulated lower amount of sRNAs (Supplementary Figure 1B). This integrative analysis also allowed us to identify genes whose mRNA levels increase during macrophage interaction while sRNA production decreases, as these genes could play a crucial role in fungal pathogenesis by being selectively protected from degradation. The total number of genes exhibiting this behaviour was 24 in WT-M vs WT-C and 127 in *r3b2*Δ-M vs *r3b2*Δ-C (Figure 3B). In addition to its possible role during pathogenesis, genes that show a pattern in which the increase in mRNA levels is associated with a reduction of sRNA levels (-sRNAs, +mRNA) were considered as direct targets of the R3B2 activity. This group includes 24 in *r3b2*Δ-M vs WT-M (Figure 3C), constituting 5 genes in the *r3b2*Δ-C vs WT-C. A functional analysis revealed that the majority of them are involved in signal transduction and signaling, intracellular trafficking, transcription, replication, and multiple metabolic processes (Supplementary Figure 1).

### Fungal genetic markers of early phagocytosis and NCRIP regulation

To identify fungal genes that serve as molecular markers of early phagocytosis or NCRIP regulation, five genes that were differentially expressed in both *r3b2*Δ-M vs *r3b2*Δ-C and WT-M vs WT-C comparisons were transcriptionally validated by RT-qPCR (Figure 3D). An LTR transposon (ID: 81899) was actively transcribed in the *r3b2*Δ mutant but downregulated during its interaction with macrophages. Interestingly, this transposon is not actively transcribed in the WT strain, either during macrophage interaction or saprophytic growth. Supporting this, sRNA analysis reveals a higher abundance of sRNAs targeting the transposon for degradation by the NCRIP in the WT strain (Figure 3E). Additionally, RT-qPCR confirms that this transposable element is consistently downregulated during macrophage interaction (in both the mutant and WT strains) and under saprophytic conditions (exclusively in the WT strain) (Figure 3D). These findings suggest that the NCRIP tightly controls LTR-transposon activity under non-stress conditions. Notably, in the absence of NCRIP activity (*r3b2*Δ-M vs *r3b2*Δ-C, Supplementary File 1), as well as during macrophage interaction in the *r3b2*Δ mutant, transposon expression decreases further (Figure 3E).

Similarly, RT-qPCR validation confirmed that the genes encoding two transcriptional regulators —a BRCT domain-containing protein (Brca1) (ID: 113714; KOG4362) and an HMG-box transcription factor (ID: 104613; KOG2746)—as well as an amino acid transporter permease (ID: 165343; KOG1286) were significantly upregulated during early phagocytosis of *M. lusitanicus* spores (Figure 3D), reinforcing their potential role in fungal adaptation to host immune responses. Conversely, the Prd transcription factor (ID: 83143; KOG0849) appears to be more relevant during saprophytic growth, as demonstrated by its expression pattern in both the RNA-seq data and RT-qPCR validation.

### Chronological transcription of chromatin and transcription-related genes during *M. lusitanicus* phagocytosis

To explore the contribution of RNA interference (RNAi) pathways and the impact of early macrophage phagocytosis on fungal gene regulation, we focused on four candidate genes showing differential expression at the small RNA (sRNA) and/or messenger RNA (mRNA) levels under various experimental conditions (WT-M, WT-C, *r3b2*Δ-M, and *r3b2*Δ-C). These genes included two chromatin-related proteins—Histone 1 (GeneID: 83400), a histone deacetylase (GeneID: 168144)—and the two above-mentioned transcription regulators— Box (GeneID: 104613) and Brca1 (GeneID: 113714) (Figure 4A).

**Fig. 4.**
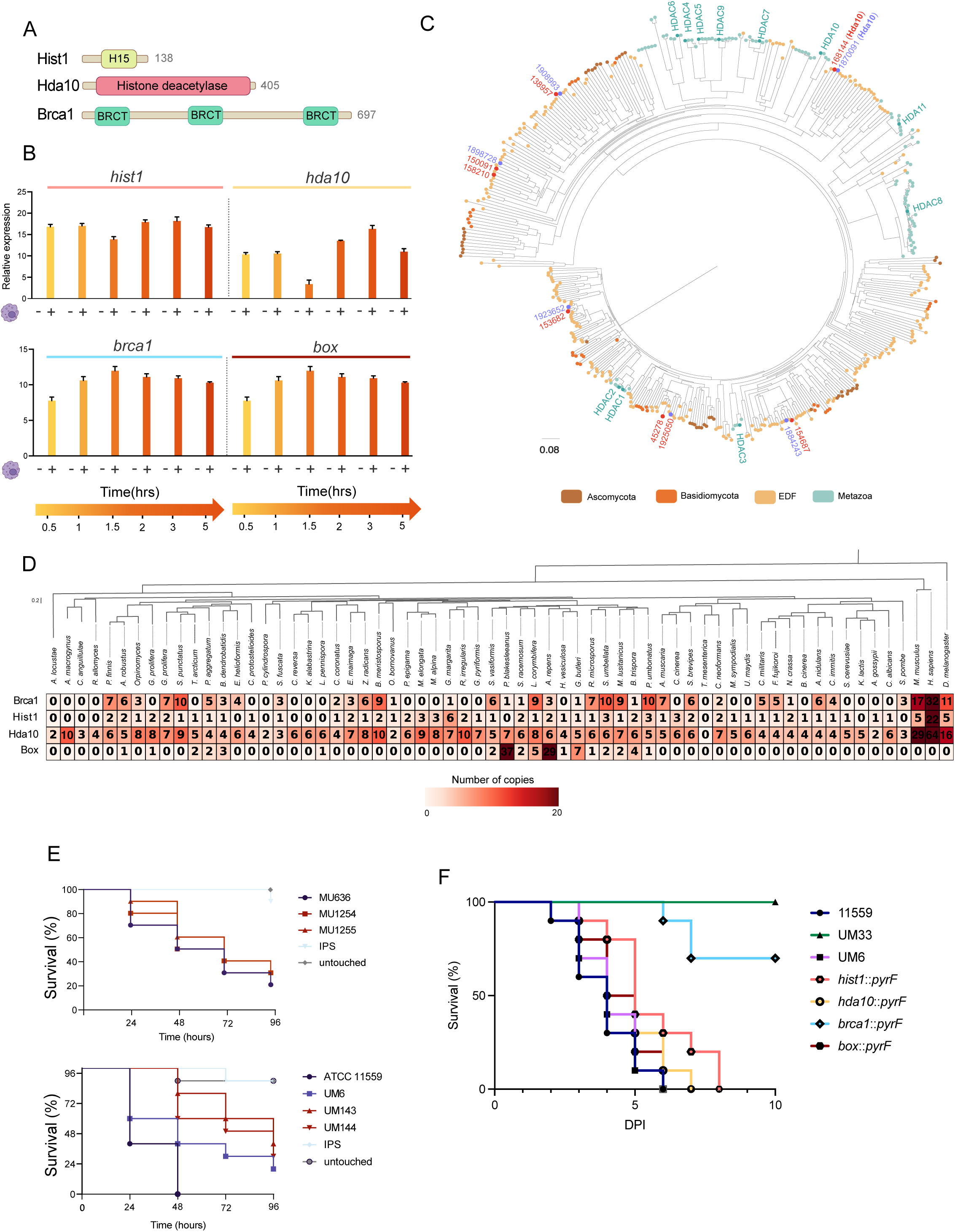
**(A)** Domain architecture of candidate virulence factor genes. Schematic representation of protein domain structures for the selected genes involved in chromatin remodelling and transcriptional regulation: the transcription factors *brca1* and *box*, and the chromatin-associated genes *hist1* and *hda10*. **(B)** Temporal expression profile of candidate virulence genes during macrophage interaction. Quantification of gene expression levels for *brca1*, *box*, *hist1*, and *hda10* in the wild-type (WT) strain of *M. lusitanicus* during phagocytosis by macrophages at 30 min, 60 min, 90 min, 2 h, 3 h, and 5 h post-infection. Gene expression in fungal spores without macrophage interaction was also evaluated by comparing WT under control conditions (WT–C) to the RNAi-defective mutant (*r3b2*Δ–C), to identify baseline regulatory differences. (C) Phylogenetic analysis of HDA homologs. Maximum likelihood phylogeny of histone deacetylase (HDA) proteins from *M. lusitanicus*, *R. microsporus* (highlighted by red dots), and other representative species from the Early Diverging Fungi (EDF), Ascomycota, Basidiomycota, and Metazoa. Human HDA isoforms were included as outgroups to classify fungal HDAs and infer potential functional conservation. **(D)** Distribution and conservation of candidate virulence genes across fungal species and Metazoa. Comparative phylogenomic analysis showing the presence/absence and number of homologs for BRCA1, Box, HDA10, and Histone 1 across multiple fungal genomes. Homolog identification was performed using InterProScan based on conserved domain architectures. Due to the lack of identifiable domains in some Box orthologs, domain-based filtering was omitted for this gene. Notably, Box appears to be conserved exclusively within fungal lineages. **(E)** Virulence assays of the transcription factors and chromatin-related mutants challenged against *G. mellonella* and murine model. The average survival rate of 3 independent survival assays in *G. mellonella* challenged with *M. lusitanicus* and *R. microsporus* spores. Statistics were done utilizing graph pad prism software. Significance was determined with log-rank (Mantel–Cox) test, utilizing GraphPad Prism. Differences were considered significant at p-values ≤ 0.05. Virulence assays where spores of the phagocytosis-related mutants in *brca1::pyrF, box:pyrF, hist1::pyrF,* and *hda1::pyrF* were injected via retroorbital into healthy mice. The survival curves of each group of 10 mice are represented by colors. A Mantel-Cox test was conducted to determine the significant differences in virulence. The avirulent control UM33 strain and the virulent UM6 have P values of P<0.0001 and P=0.644, respectively. DPI are days post-injection.

The genes encoding histone deacetylase and Brca1 exhibited a strong accumulation of both sRNA and mRNA in the *r3b2*Δ mutant during macrophage interaction (*r3b2*Δ-M vs *r3b2*Δ-C), suggesting regulation by the canonical RNAi pathway and additional induction by phagocytic stress (Supplementary Figure 2). In contrast, *box* and *his1* displayed expression changes mainly at the mRNA level, with minimal sRNA variation, implying RNAi-independent transcriptional regulation.

Next, since phagocytosis of *M. lusitanicus* spores leads to transcriptional reprogramming, likely involving chromatin remodelling and transcriptional regulation necessary for survival and germination within the phagosome total RNA was extracted from the WT strain at various time points: 30 min, 60 min, 90 min, 2 h, 3 h, and 5 h, as well as from L15 medium as a non-infection control. RT-qPCR analysis revealed that the chromatin component *hist1*, the chromatin modifier histone deacetylase, and both transcriptional regulators were highly expressed during the phagocytic process compared to non-stressful conditions (Figure 4B). These genes, which are involved in regulating downstream gene expression, suggest that they might play a key role in controlling the expression of virulence-related genes.

### Functional characterization of the RNAi-regulated and phagocytosis marker genes

To better understand the biological implications and evolutionary significance of these genes, we performed a comprehensive conservation analysis. Using *M. lusitanicus* as the reference organism, we examined the presence and evolutionary conservation of these proteins in 61 species from major fungal phyla. Additionally, we expanded the analysis to include metazoan species, such as *Mus musculus*, *Homo sapiens*, and *Drosophila melanogaster.* In the case of histone deacetylases (HDAs), we constructed a phylogenetic tree using the complete set of well-characterized human HDAs as references. In *M. lusitanicus*, we identified seven distinct HDA-encoding genes, and we found that the gene studied in this work encoded for a putative ortholog of the human HDA10, supporting its annotation as *hda10* (Figure 4C). Domain-based searches of proteins containing BRCT (for *brca1)* revealed substantial variability in the number of such domain-containing proteins across species. For *brca1*-related proteins, which include BRCT domain-containing factors involved in DNA repair, transcriptional regulation, and cell cycle control (13), the number of detected proteins ranged from 0 (e.g., *H. vesiculosa*, *U. maydis*) to 32 in *Homo sapiens*, suggesting both gene duplication events and potential gene losses throughout evolution (Figure 4D). Similarly, histone H1 domain-containing proteins—key players in DNA compaction and chromatin structure (14)—ranged from 1 to 6 copies in several fungal species, while others, such as *P. blakesleeanus*, lacked detectable homologs. Interestingly, both *M. lusitanicus* and *R. microsporus* possess only a single copy of histone H1, which may reflect a more streamlined chromatin organization or reduced redundancy in these species(Figure 4D).

Proteins containing histone deacetylase (HDA) domains also exhibited significant variation in copy number. Species such as *Rhizopus irregularis*, *R. microsporus*, and *M. lusitanicus* contained between 6 and 10 copies (Figure 4C and Figure 4D), while others, including *A. locustae* and *S. pombe*, had only 2 to 3 copies. In contrast, metazoan species showed much higher copy numbers, with *Homo sapiens* having 64 copies of proteins containing the histone deacetylase domain, *Mus musculus* with 29, and *Drosophila melanogaster* with 16 copies. Interestingly, *box* homologs were found exclusively in fungi, with no representatives detected in the metazoan genomes analyzed (Figure 4D).

### Disruption of the chromatin and transcription-related genes impairs *R. microsporus* virulence and and fungal fitness, and suggests a trend toward attenuation in *M. lusitanicus*

To assess their role in virulence and fungal development, and to elucidate the interplay between canonical and non-canonical RNAi pathways in the context of host-pathogen interactions, we generated single deletion mutants in *M. lusitanicus* (Supplementary Figure 4A). To evaluate whether these genes perform conserved functions across species, the corresponding orthologs in *R. microsporus* were also disrupted (Supplementary Figure 4B).

First, the mutant strains were phenotypically characterized by monitoring their sporulation (Supplementary Figure 4A, C, D, G) and vegetative growth (Supplementary Figure 4B, E). *hist1* mutant in *R. microsporus* presented a lower growth rate (Supplementary Figure 4F) and also reduced sporulation (Supplementary Figure 4D, G). However, neither the *hist1*Δ in *M. lusitanicus* nor the rest of the mutants, either in *M. lusitanicus* or in *R. microsporus*, showed defects in these biological processes.

Additionally, the effects of temperature and light on mutant growth were evaluated under rich medium conditions. Experiments conducted in darkness revealed that only the *brca1* mutant of *R. microsporus* exhibited a notable phenotype, showing increased sporulation at 37 °C compared to the wild-type strain (Supplementary Figure 5). This observation suggests that disruption of this transcription factor induces a stress response specifically under elevated temperature conditions.

Subsequently, a first screening of their impact on pathogenicity was performed using the *Galleria mellonella* infection model (Figure 4E, Supplementary Figure 6), allowing us to evaluate their contribution to virulence in both species. *G. mellonella* larvae were challenged with the *M. lusitanicus* and *R. microsporus* strains (Supplementary Table 1). In *M. lusitanicus*, mutants lacking *brca1*, *box*, and *hda10* showed no significant differences in virulence between independently generated deletion strains. However, in the case of *hist1Δ*, mutant MU1248 displayed a consistent trend toward reduced virulence compared to its replicate strain and the wild-type. Similarly, mutant MU1250 (*brca1Δ*) also exhibited lower survival rates in infected hosts. Although variability across biological replicates limited statistical significance, these findings suggest that deletion of *hist1* and *brca1* may negatively impact virulence in *M. lusitanicus* (Figure 4E, Supplementary Figure 6A). In contrast, more robust and statistically supported phenotypes were observed in *R. microsporus*. Deletion of *brca1* (*brca1::pyrF*) resulted in significantly attenuated virulence in both independent mutant strains (UM141 and UM142), supporting a conserved role in pathogenicity (Supplementary Figure 6B). Similarly, disruption of *box* (*box::pyrF*) led to reduced virulence: both UM144 and UM143 showed increased host survival, with UM144 displaying statistically significant attenuation compared to both the wild-type (ATCC 11559) and control strain UM6 (Figure 4E, Supplementary Figure 6B).

Mutants lacking *hda1* or *hist1* presented more variable outcomes. For *hda1::pyrF*, differences between mutant strains were noted, yet one isolate (UM121) exhibited significantly reduced virulence compared to the wild-type. In the case of *hist1::pyrF* mutants, although no significant differences were observed relative to the control strain UM6, both mutants showed markedly lower mortality compared to the wild-type, suggesting a potential, albeit modest, impact on virulence. Additionally, to study if the disrupted genes in *R. microsporus* influence the fungal pathogenesis, virulence assays were conducted using mice. The UM141 (*brca1::pyrF)* strain, was the most affected mutant (P<0.0001) in virulence (Figure 4F), showing almost an avirulent phenotype, suggesting a crucial implication of Brca1 in promoting fungal virulence during infection. On the other hand, the chromatin-related mutants, *hist1::pyrF* (UM145 strain) and *hda1::pyrF* (UM121), show impairments in virulence (P=0.0204 for *hist1::pyrF*, P=0.0372 for *hda10::pyrF*) (Figure 4F), causing pathogenicity attenuation. Finally, the UM143 (*box::pyrF,* P=0.2317) strain did not display significant effects on the fungal virulence in healthy mice.

## DISCUSSION

In response to fungal invasion, macrophages act as the first line of defense, recognizing and internalizing Mucoralean spores to initiate phagosome maturation. This process exposes the spores to acidic pH, reactive oxygen species, and nutrient deprivation, aiming to eliminate them (4). However, depending on the fungal species and the immune status of the host, some spores can survive, germinate, and escape, leading to infection (4). While previous studies have shed light on the later stages of infection (12), the genetic regulation underlying the early interaction between *M. lusitanicus* and macrophages remains largely unexplored. To address this gap, we selected *M. lusitanicus* as a model organism due to its genetic tractability and included *R. microsporus* —a major causative agent of mucormycosis— due to recent advancements in its genetic manipulation (15). This dual-species approach aimed to identify conserved virulence factors and unravel fungal strategies that facilitate immune evasion and pathogenesis.

In this work, we provided new insights into the transcriptional landscape of early phagocytosis in Mucorales by examining the internalization of two *M. lusitanicus* strains by macrophages: a virulent wild-type and an RNAi-deficient mutant. The transcriptomic analysis revealed thousands of differentially expressed genes (DEGs), with notable upregulation of genes involved in amino acid transport, energy metabolism, and translation, suggesting immediate metabolic adaptation to phagosome stress. This metabolic activation is consistent with findings from previous studies on the late stages of *M. lusitanicus* phagocytosis (16). Together, these results contribute to our understanding of the adaptive response of the pathogen to host-mediated immune challenges, further emphasizing the importance of these early transcriptional shifts in fungal survival and pathogenesis.

Interestingly, the comparative transcriptomic response during the early and late phagocytosis of *M. lusitanicus* spores indicates that the fungus mounts a major transcriptional response in the early stages of infection, which prompted us to further analyze the chronological sequence of the host-pathogen interaction. While many of the DEGs were common to both stages, we also identified DEGs unique to the early phase of infection, likely involved in recognition, adhesion, and the transcriptional response necessary to initiate fungal germination. Additionally, we identified exclusive genes associated with the late phagocytosis phase, suggesting their involvement in immune evasion mechanisms. These findings emphasize the dynamic fungal response to the host immune system, highlighting the importance of both early and late phases in the infection process.

In our study, we integrated the analyses of sRNA and mRNA profiles during the early phagocytosis of *M. lusitanicus* spores by macrophages to understand gene regulation. The data showed that the the *r3b2*ΔM mutant had significantly more differentially degraded genes (DDGs) during macrophage interaction, compared to the WT strain, suggesting activation of the canonical RNAi pathway in absence of NCRIP. Additionally, sRNA production was higher in the WT strain relative to the mutant, both during infection and under saprophytic growth, supporting the idea that NCRIP is is a key regulator of gene expression. We also identified genes whose mRNA levels increased while sRNA levels decreased, suggesting selective stabilization of these genes, which could play a role in immune evasion and fungal survival inside the host. These genes are involved in critical processes such as signal transduction, intracellular trafficking, transcription, replication, and metabolism, highlighting their potential importance in fungal pathogenesis.

The transcriptional alterations observed in *M. lusitanicus* spores during phagocytosis suggest a coordinated need for chromatin remodelling and transcriptional reprogramming for survival and germination within the host phagosome. In this context, we focused on two chromatin-related genes—*histone 1* (*hist1*) and *histone deacetylase 1*0 (*hda10*)—as well as two transcription factors—*BReast CAncer gene 1* (*brca1*) and *HMG-BOX* (*box*)— due to their putative roles in orchestrating gene expression programs linked to virulence. Our findings suggest that these genes may be regulated by both transcriptional and RNA interference (RNAi) mechanisms. Specifically, *hda10* and *brca1* showed increased accumulation of both small RNAs and mRNA under macrophage-induced stress, indicating RNAi regulation. In contrast, box and hist1 were mainly regulated at the transcriptional level, suggesting they respond to infection signals via transcription factor networks or chromatin changes independent of RNAi. These results highlight a model where *M. lusitanicus* integrates RNAi, transcriptional, and chromatin-based mechanisms to regulate gene expression and contribute to fungal adaptation and virulence during host interaction.

We deleted these four candidate genes in *M. lusitanicus* and disrupted them in *R. microsporus* to identify conserved virulence factors across Mucorales. Among these, the brca1 ortholog exhibited a notable phenotype. Disruption of *brca1* in *R. microsporus* led to a significant reduction in virulence in a murine model. Deletion of the same gene in *M. lusitanicus* also resulted in reduced virulence in the *G. mellonella* model, highlighting a previously unrecognized role for *brca1* in fungal pathogenicity. This finding suggests that the role of BRCA1 in DNA repair and genome stability (13) extends to its contribution to infection, helping fungi withstand host-induced genotoxic stress. Interestingly, our phylogenomic analysis showed variability in BRCA1 domain-containing proteins across species, with some fungi lacking detectable orthologs while others had multiple gene copies. This suggests that BRCA1-related proteins may have evolved differently across species, potentially driven by host interaction or environmental adaptation pressures. A similar pattern was observed for *histone 1*. Despite its non-essentiality for viability— consistent with previous findings in other fungi such as *S. cerevisiae* and *N. crassa* (17)— plays a role in virulence. The *hist1::pyrF* mutant of *R. microsporus* showed significantly reduced virulence in a murine model, along with growth and sporulation defects. In contrast, deletion of *hist1* in *M. lusitanicus* resulted in a trend toward reduced pathogenicity in *G. mellonella*, further highlighting species-specific differences. Phylogenomic analysis confirmed that most fungal species analyzed carry only one or two copies of *histone 1* with *M. lusitanicus* and *R. microsporus* having a single copy. These findings indicate that histone H1 has a conserved but non-essential role in fungal viability and suggest its involvement in virulence, potentially through chromatin-mediated transcriptional regulation during host interaction.

A comparable trend was observed for the histone deacetylase *hda10*, which was upregulated and showed a strong accumulation of small RNAs during macrophage interaction, particularly in the *r3b2*Δ mutant. This suggests regulation by the RNAi machinery and induction by phagocytosis-related stress signals. Conservation analysis confirmed the widespread presence of Hda10 orthologs across diverse fungi and classified it as an Hda10-type histone deacetylase. Functional studies revealed that deletion of *hda10* in *R. microsporus* (*hda10::pyrF*) resulted in reduced virulence in the murine model, while deletion of the corresponding gene in *M. lusitanicus* did not significantly impact virulence in the *G. mellonella* model. These findings are consistent with prior reports in human fungal pathogens such as *Candida albicans*, *C. glabrata*, and *Cryptococcus neoformans*, where HDACs regulate key virulence factors, including morphogenesis, stress response, and drug resistance, highlighting their potential as antifungal targets (18) (19). Together, our results suggest that Hda10 protein contributes to fungal pathogenicity in *R. microsporus* and, potentially, in other Mucorales, likely through chromatin-mediated transcriptional responses activated during host interaction. Finally, the deletion of the *Box* gene, which encodes a transcription factor, also exhibited notable effects on virulence in *R. microsporus*. Specifically, the *box::pyrF* mutant showed reduced virulence in the *G. mellonella* infection model, indicating its role in modulating pathogenicity. However, no reduction in virulence was observed in the murine infection model, suggesting species-specific differences in its influence on virulence. Notably, our conservation analysis revealed that *Box* is primarily conserved among fungi, highlighting its specialized role in fungal pathogenesis and its absence in a broad range of eukaryotes. This specificity suggests that *Box* could be a novel target for antifungal therapies, particularly in pathogens like *R. microsporus*.

Altogether, these virulence and phenotypic analyses indicate that while *M. lusitanicus* may be as a useful model organism for genetic manipulation, it may not be the best model for studying Mucorales pathogenesis, particularly regarding virulence mechanisms. Other species, such as *R. microsporus*, might provide a more accurate representation of the virulence factors at play in these fungi. Furthermore, although *M. lusitanicus* showed reduced virulence in certain mutants, it is important to note that in immunocompromised hosts, *M. lusitanicus* retains its virulence potential (12). These findings emphasize that different Mucorales species likely possess distinct virulence determinants, suggesting that a combination of model systems is essential for a comprehensive understanding of the molecular mechanisms driving fungal infections.

## ACKNOWLEDGMENTS

This research was supported by the grant PID2021-124674NB-I00 to F.E.N. and V.G, funded by MICIU/AEI/10.13039/501100011033 and by ERDF/EU.

## AUTHOR CONTRIBUTIONS

G.T. conducted most of the *M. lusitanicus* and *R. microsporus* experiments, performed the host-pathogen interactions with macrophages, prepared the *M. lusitanicus* material for sequencing, performed bioinformatic analysis of the results, prepared the figures and tables, designed and coordinated the project, and wrote the manuscript with significant input from C.L., F.E.N. and V.G. C.L participated in the generation of *M. lusitanicus* and *R. microsporus* mutant strains, in the bioinformatic data processing, results representation and virulence assay conducted with mice. U.B. and J.S. performed the virulence assays with *Galleria mellonella.* E.N. provided materials. F.E.N. supervised and coordinated the project. V.G. supervised and coordinated the project.

## METHODS

### Fungal strains and culture conditions

The *M. lusitanicus* strains MU636 (*leuA^-^*) and MU412 (*leuA, r3b2-^-^*) were used in the host-pathogen in vitro interaction. Both strains have the same genetic background (20) (21). MU636 was employed as a recipient strain for the deletion of the transcription regulators *brca1* and *box* and the chromatin related genes *hist1* and *hda10*. Also, together with R7B, MU636 was used in the *Galleria mellonella* infection. On the other hand, the NRRL3631 was the avirulent control.

For optimal growth and sporulation, *M. lusitanicus* strains were grown in rich medium (YPG; 3 g/L yeast extract, 10 g/L peptone, 20 g/L glucose, 15 g/L agar) pH 4.5 at 26°C and under illumination. Subsequently, spores were harvested, washed, and filtered using 70μm strainer cells before the host-pathogen interaction.

All the *R. microsporus* strains used in this work derived from the WT *Rhizopus microsporus* ATCC 11559 (listed in Supplementary Table 1), which was used in the mouse infection as the virulent control. UM6 was also used as a virulent strain. The avirulent control was the UM33, which was employed as the receptor strain for the mutant generation. All *R. microsporus* strains were cultured in rich medium YPG pH 4.5 at 30°C for spore harvesting, supplemented with uridine (200 mg/l) when required. All the media and culture conditions used for *M. lusitanicus* and *R. microsporus* growth are detailed in Lax et al. (15).

### In vitro host-pathogen interactions

The in vitro interaction of MU412 and MU636 spores with macrophages was conducted following the procedure described in Pérez-Arques et al. (16). Briefly, *M. lusitanicus* spores were confronted with the mouse cell line of macrophages J774A.1; ATCCTIB-67 in a proportion of 1.5:1 (spores: macrophages). For the RNA-seq analysis, spores were exposed to macrophages for 1 h to conduct early phagocytosis. For the chronological expression of the gene candidates involved in the early phagocytosis for the q-RT-PCR analysis, confrontations of MU412 and MU636 spores to macrophages were performed for 30 min, 60 min, 90 min, 2 h, 3 h, and 5 h. The in vitro interaction with macrophages was performed at 37°C in an L15 medium (Capricorn Scientific) supplemented with Fetal Bovine Serum (FBS, Capricorn Scientific). For the saprophytic conditions, the same concentration of MU412 and MU636 spores was cultured in L15 supplemented with FBS. For the Lysotracker Green DND-26 (ThermoFisher) and DAPI (Life Technologies) staining, the protocol described in (7) was followed.

### RNA extraction, library preparation, and RNA-seq analysis

For the in vitro host-pathogen interactions, total RNA was extracted from three replicates of MU636 and MU412 strains interacting with macrophages (after 1 h) and three replicates of each of these strains growing for 1 h in saprophytic conditions. Total RNA (sRNA + mRNA) was purified from the 12 samples using the MirVana Kit (InvitrogenTM) and following the procedure for sRNA enrichment. Each sample was divided into two, one for mRNA sequencing and the other for sRNA sequencing. Samples were quality-checked, quantified, and deeply sequenced by Novogene.

For the in vivo infection, three replicates from the host-pathogen interaction were used for total RNA extraction using the mirVana kit. Additionally, three replicates of the fungal spores growing in saprophytic conditions, as well as three replicates from the mouse cell samples (in a non-infection context) were also extracted and deeply sequenced by the same company. Raw sRNA-seq reads were quality-checked with FASTQC v0.11.8 (https://www.bioinformatics.babraham.ac.uk/projects/fastqc/) before and after removing adapter (3´) and contaminant sequences and low-quality reads (-q28-p50) were removed with Cutadapt v3.4 (https://cutadapt.readthedocs.io/en/v3.4/). Clean reads between 18 and 35nt were aligned to the *M. lusitanicus* v2.0 genome (retrieved from https://mycocosm.jgi.doe.gov/Mucci2/Mucci2.home.html) (22) using Bowtie2 v2.5.3 (23). Reads mapping on the *M. lusitanicus* genome were analysed together with FeatureCounts v2.0.1 (24). The software produced a count table file (with several reads from each library that defined each locus), which was used for differential expression (DE) analysis between the mutant and wild-type strains (with or without macrophages for the in vitro host-pathogen interaction) with DESeq2 v2.11.40.6 (25). Genes with an adjusted p-value ≤0.05 and a log2 fold-change |(log2FC) ≥1| were considered as differentially expressed.

Raw mRNA datasets were checked for quality with FASTQC before and after removing adapter and contaminant sequences with TrimGalore! v0.6.2 (https://github.com/FelixKrueger/TrimGalore) excluding reads with a Phred quality score Q≤32 and/or a total length ≤20nt as well as adapter sequences with an overlap ≥4 bases. The mRNA clean reads were aligned to the *M. lusitanicus* v2.0 genome using HISAT2 aligner (26). HTSeq software v.2.0.5 (27) was used to quantify gene expression levels, and differential expression analysis was performed using DESeq2, considering genes with an FDR ≤ 0.05 and |log2FC| ≥ 1 as significantly differentially expressed.

KOG class enrichment analyses for *M. lusitanicus* DEGs were performed in GraphPad Prism v8.0.1; delta-ranks were computed as the difference between the mean of all genes within the KOG class and the mean rank of all other genes in a Mann-Whitney U-test. A KOG class was considered over-represented if P≤0.05 in one-sided Fisheŕs exact evaluation (performed in R) of the DEGs compared to the total amount of genes in each KOG class.

### q-RT-PCR quantification

Once total RNA was extracted, about 5μg of total RNA of *M. lusitanicus* WT (MU636) and mutant (MU412) strains samples were treated with the Turbo DNase (Thermo Fisher). The RNA samples were routinely checked for DNA contamination by a PCR analysis using primers for the housekeeping elongation factor gene. For cDNA synthesis, 1μg of total RNA was reverse-transcribed using the iScript cDNA synthesis kit (Bio-rad) at 25°C for 10min, 42°C for 50min, and 70°C for 15min. The RT-qPCR was performed in triplicate using 5X SYBER green PCR master mix (Applied Biosystems) with a QuantStudio TM 5 flex system (Applied Biosystems) following the supplieŕs recommendations. To ensure non-specific amplification, non-template control and melting curves were tested. The primer sequences used for the quantification of genes are listed in Supplementary Table 2. The efficiencies of every pair of primers were approximately identical, and the relative expression of target genes was obtained by the delta-delta cycle threshold (ΔΔCt) method, normalizing for the endogenous control elongation factor gene (*ef*).

### Nucleic acid manipulation and plasmid constructions

All PCR amplifications were conducted using the Herculase II fusion DNA polymerase (Agilent, Santa Clara, California, USA), with annealing temperature and extension time adjusted based on the specific pair of primers used and the fragment length to be amplified, respectively. PCR fragments were purified from gel when required using the GeneJet Gel Extraction kit (Thermo Scientific). DNA extraction was conducted following the previously described protocol in (21). All primers were designed using the Primer3Stat (https://www.bioinformatics.org/sms2/pcr_primer_stats.html) and Multiple Primer Analyzer (https://www.thermofisher.com/es/es/home/brands/thermo-scientific/molecular-biology/molecular-biology-learning-center/molecular-biology-resource-library/thermo-scientific-web-tools/multiple-primer-analyzer.html) tools.

### Deletion cassettes for phagocytosis-related mutants in *M. lusitanicus*

The phagocytosis-related single mutants were achieved using genetic constructions consisting of the selectable marker *leuA* flanked by 1Kb upstream and downstream of the gene candidates. The 1Kb upstream, the 1Kb downstream, and the selectable marker were PCR amplified using specific primers (Supplementary Table 2). Subsequently, by overlap PCR the three fragments were fused generating the deletion fragment. The deletion cassettes were purified and used for the protoplast transformation of the MU636 *M. lusitanicus* strain. Transformants were acquired following the previous transformation procedure (28). In short, harvested fresh spores were incubated for 2-4 hours ensuring their correct germination. Subsequently, protoplasts were obtained by cell wall digestion of the germinated spores with lysing enzymes (Sigma-Aldrich) and chitosanase (Sigma-Aldrich). The protoplast transformation was performed by electroporation with *SaclI* linearized plasmids and later incubated in poor agar medium YNB + Sorbitol 3.2 and checked 3-4 days for colonies. Approximately, 5μg of of overlapping PCR products, which include the *leuA* gene flanked by 1Kb upstream and downstream the open reading frame (ORF) of each of the candidate genes, was necessary for each transformation. As spores are multinucleated, colonies were subsequently transferred to fresh YNB agar plates (5-10 vegetative cycles) to obtain homokaryonts with all nuclei transformed. DNA from transformants was extracted as previously described. Homokaryosis was checked using primers that bind upstream and downstream the 1Kb employed for the fusion PCR to ensure the absence of WT nuclei (Supplementary Figure 4).

### Disruption cassettes for phagocytosis genes in *R. microsporus*

Disruption of the orthologous genes in *R. microsporus* (*hist1* [ID: 1913394], *hda10* [ID: 1870091], *brca1* [ID: 1897364], and *box* [ID: 1900053]) was performed using the CRISPR-Cas9 system, following the genetic disruption protocol described by Lax et al. (29). Briefly, crRNAs (Supplementary Tablev3) were designed using the EuPaGDT gcRNA design tool (http://grna.ctegd.uga.edu/) with default parameters to guide the Cas9 cut. For homology-directed repair, primers amplifying the *pyrF* selective marker were designed with 38 bp of homology upstream and downstream of the cut point. To verify integration, a reverse primer specific to *pyrF* and a forward primer that hybridizes upstream of the cut site were designed. Homokaryon checking was performed using the forward upstream primer and a reverse downstream primer to the cut site. All primers were designed using Primer3Stat and the Multiple Primer Analyzer tools.

For the transformation of *R. microsporus*, the UM33 recipient strain (*pyrF-*, *LeuA-*) was used. Protoplasts of UM33 were transformed by electroporation with 150–200 mg/ml of linear DNA fragment (pyrF flanked by 38 bp tails). Transformed protoplasts were then cultured on MMC selective media (without uridine). Genomic DNA of the transformants was extracted following the procedure outlined by Osorio-Concepcion et al. (21) for integration and homokaryon checking.

### Phenotypic characterization of the phagocytosis mutants

Radial growth and sporulation of the phagocytosis mutants were analysed, inoculating drops of 500 spores in the plate center of MMC 3.2 (for radial growth) and YPG 4.5 (for sporulation). Radial growth was monitored during 5 d each 24 h through measuring the colony diameter. Spore production was determined by quantifying the spore concentration of 1cm^2^ of a chunk of agar, which was later transferred to a 50 ml falcon containing 10ml of PBS1X. Subsequently, the falcon was vigorously vortexed to enable spore release, which were counted using a Neubauer camera, determining the spore concentration of the mutant and the WT strains. Both radial growth and spore production were quantified using three replicates per mutant. Statistical analysis was conducted using one-way ANOVA followed by Student’s *t*-test, considering differences statistically significant at *p* ≤ 0.05.

### Virulence assays

To determine if the phagocytosis-related genes constitute conserved virulence factors in Mucorales, as a first screening, *G. mellonella* larvae were infected with *M. lusitanicus* mutants and the corresponding ortholog mutants in *R. microsporus.* To this end, sixth instar larvae of *G. mellonella* (SAGIP, Italy), weighing 0.3 -0.4 g, were selected for experimental use. Larvae, in groups of twenty, were injected through the last pro-leg into the hemocoel with 106 spores in a volume of 20 μl according to Kelly & Kavanagh (30) and incubated at 37°C. Untouched larvae and larvae injected with sterile insect physiological saline (IPS) served as controls. Survival was monitored every 24 h for the duration of 96 h. Experiments were repeated at least 3 times.

On the other hand, to assess the role of phagocytosis-related genes in *R. microsporus* virulence, survival assays were conducted as previously described by Lax et al. (15). *M. lusitanicus* mutants were discarded from the mice infection, due to the absence of significant reduction in virulence obtained in *G. mellonella.* Briefly, healthy Swiss mice weighing ≥30 g and one month old (supplied by the Animal Facility Services, University of Murcia, Spain) were used as the host model for virulence assays. Groups of 10 mice (per strain) were injected via retroorbital infection with suspensions of 1.5×10^6^ from the mutant or the WT strain. Mice were housed under established conditions with free food access and sterile water. The survival rates of each group were checked twice a day for 15d, and mice reaching the endpoint were euthanized in a CO^2^ chamber. These rates were represented in a Kaplan-Meier curve (GraphPad Prism). Both in *G. mellonella* and in mice virulence assays, statical analysis was performed by Mantel–Cox test, and differences in survival were considered significant with a P value ≤ 0.05.

### Ethics statement

The virulence assays in this work were performed according to the ethical Guidelines of the European Council (Directive 2010/63/EU) and the Spanish RD 53/2013, ensuring animal welfare, by minimising any potential pain, distress, or suffering experienced by the mice along the course of the experiments. Procedures and experiments conducted in this study were closely supervised and granted approval by the University of Murcia Animal Welfare and Ethics Committee, as well as the Council of Water, Agriculture, Farming, Fishing and Environment of Murcia, Spain (authorization number REGA ES300305440012).

### Phylogenetic analysis and ortholog search

Proteomes of 64 representative species were retrieved from the Joint Genome Institute (JGI) MycoCosm genome portal (31) and Uniprot (32). Sequences of *M. lusitanicus* Histone 1 (ID: 83400), Hda10 (ID: 168144), Brca1 (ID:113714), Box (ID: 104613) proteins were queried against the selected proteomes using iterative HMMER jackhmmer searches (E-value ≤1×10-3) (v3.3.2) (http://hmmer.org/). A reciprocal BLASTp search (v2.10.1) (33) was conducted and sequences that failed to produce a hit were discarded. An additional search using Pfam-A database (34) using HMMER hmmscan (v3.3.2) (http://hmmer.org) served to remove hits that lacked the BRCT domain and BRCA1-associated (Brca1 search), the Histone deacetylase domain (HDA search), or the Linker histone H1/H5, domain H15 (Histone 1 search) (Supplementary File 3). For the Box search, domain-based filtering was not applied, since most Box orthologs identified in fungi do not contain any known conserved domains. The remaining Histone 1, Hda10, BRCA1, and Box candidate sequences were used to determine the copy number of each protein type across all analyzed species. This data was compiled into a matrix representing the number of copies per species, which was subsequently used to generate a heatmap illustrating the distribution and expansion of each protein family across fungal lineages. The Hda candidate sequences were further aligned using Clustal Omega (35) with default parameters. A phylogenetic tree was then generated from this alignment using Clustal Omega’s built-in approximate maximum-likelihood method with default settings. The species tree was generated after analyzing the 64 proteomes with OrthoFinder (Inflation factor, -I 1.5) (36). Phylogenetic trees were visualized in iTOL (37).

## DATA AVAILABILITY

The raw sequence data that support the findings of this study have been deposited in the Sequence Read Archive (SRA) under the accession numbers and in the Gene Expression Omnibus (GEO)

## CODE AVAILABILITY

The scripts used for the bioinformatic analysis are available at https://github.com/ghizlanetahiri95/Early_Phagocytosis_Mucorales_RNAi.

**Fig. Supp. 1 (A)** Functional analysis of the genes that have an increase in their mRNA production and a reduction of their corresponding sRNAs during their interaction with macrophages (WT-M / WT-C, *r3b2Δ-*M / WT-M), and those genes that experienced these changes in the comparison of the mutant versus WT during their phagocytosis. **(B)** Correlation between sRNA production and mRNA expression levels. The top 100 genes with the highest and lowest sRNA production were selected for the WT strain with and without macrophages, and the *r3b2*Δ with and without macrophages. The mRNA expression levels of these top 100 genes, represented as the log2 of FPKM, were plotted for each group. A negative correlation is observed, where genes with higher sRNA accumulation exhibit lower mRNA expression levels, while genes with lower sRNA accumulation tend to have higher mRNA expression levels. Boxplots indicate the median, first, and third quartile; whiskers extend to the 10th and 90th percentiles. Statistical significance was assessed using Welch’s t-test. The p-values obtained were 0.0008 for WT-C, <0.0001 for WT-M, 0.03223 for *r3b2*Δ-C, and ns (not significant) for *r3b2*Δ-M.

**Fig. Supp. 2** Genomic coverage of sRNA and mRNA reads mapped to the selected genes involved in chromatin remodelling and transcriptional regulation: the transcription factors *brca1* and *box*, and the chromatin-associated genes *hist1* and *hda10* in WT and *r3b2*Δ strains, under both saprophytic and macrophage-interacting conditions. Yellow and red plots represent sRNA and mRNA read coverage, respectively.

**Fig. Supp. 3 (A)** Homokaryosis PCRs of the mutants in the potential virulence factors in *M. lusitanicus*. PCRs to check if the mutants are homokaryonts were conducted using a forward and a reverse primers that bind upstream and downstream the regions used for the fusion (Locus_F and Locus_R). The PCR products in the deleted mutant result in 5Kb, 5Kb, 5.1Kb, and 5.3Kb in the *hist1, brca1, hda10, and box* loci respectively. The PCR products with the wild-type genes have 0.63Kb, 4.8Kb, 3.5Kb, and 4.3Kb for *hist1, brca1, hda10*, respectively. Red boxes show the expected PCR products in the deletion nuclei. The heterokaryons were submitted to further vegetative cycles in selective media. PCRs to check homokaryosis of *R. microsporus* mutants in phagocytosis-related genes**. (B)** PCRs to check *R. microsporus* homokaryosis of the gene candidate disruptions, using specific primers that hybrid approximately 1Kb from the 38bp homology regions. The amplification fragments lengths (shown in red boxes) of the disrupted genes are 5.5Kb for *hist1, hda10, and brca1* disruptions, and 5.3Kb for the *box* disruption. The amplification product lengths resulting from the WT nuclei are 2Kb for the *hist1, hda10, and brca1* genes, and 1.8Kb for the *box* locus. The mutants containing only nuclei with the disruptions (red rows) were selected for the subsequent analysis.

**Fig. Supp. 4** Phenotypic characterization of the transcription regulators and chromatin-related mutants. **(A)** Sporulation of mutants with disruptions in transcriptional regulator and chromatin-related genes in *M. lusitanicus*. **(B)** Monitoring of radial growth of *M. lusitanicus* mutants every 24 hours over 5 days. **(C)** Spore production per cm² for in the WT strain and each mutant in *M. lusitanicus* after 2 days of growth. **(D)** Sporulation of the corresponding mutants in *R. microsporus*. **(E)** Monitoring of radial growth of these mutants in *R. microsporus* every 24 hours over 5 days. **(F)** Spore production per cm² for in the WT strain and each mutant in *R. microsporus* after 2 days of growth. Statistical significance was first assessed using one-way ANOVA to determine overall differences among strains. Post hoc pairwise comparisons between each mutant and the WT were then conducted using Student’s *t*-test. The *hist1-*mutant in *R. microsporus* generated a lower production of spores by cm^2^ and had a lower growth rate compared to the WT strain. In the two cases, differences were statistically significant. For radial growth, the results for hist1 mutant 1 (UM145) compared to the WT were as follows: Day 1: p-value = 0.02482, Day 2: p-value = 0.4226, Day 3: p-value = 0.005651, Day 4: p-value = 0.03553, and Day 5: p-value = 0.0078. For hist1 mutant 2 (UM146) compared to the WT, the p-values were: Day 1: p-value = 0.008388, Day 2: p-value = 0.2697, Day 3: p-value = 0.006051, Day 4: p-value = 0.01623, and Day 5: p-value = 0.003049. In terms of sporulation, mutant 1 (UM145) showed a p-value of 0.02719, while mutant 2 (UM146) had a p-value of 0.04289.

**Fig. Supp. 5** Effects of temperature and light on mutant sporulation. **(A)** Sporulation of two *R. microsporus* mutants disrupted in the *brca1* gene was evaluated under dark conditions at 37°C. The *brca1::pyrF* mutant exhibited increased sporulation compared to the wild-type strain, suggesting a stress response triggered by elevated temperature. **(B)** Plates showing sporulation differences between the mutants and WT strain at 37°C and in darkness. Statistical significance was initially evaluated using one-way ANOVA to assess overall differences among the strains. Subsequent post hoc pairwise comparisons between each mutant and the WT strain were performed using Student’s t-test. For *brca1* mutant 1 (UM141), the p-value was 0.005311, and for *brca1* mutant 2 (UM142), the p-value was 0.03884.

**Fig. Supp. 6** Virulence assays of the transcription factors and chromatin-related mutants challenged against *G. mellonella and* murine model. The average survival rate of 3 independent survival assays in *G. mellonella* challenged with *M. lusitanicus* **(A)** and *R. microsporus* **(B)** spores. Statistics were done utilizing graph pad prism software. Significance was determined with log-rank (Mantel–Cox) test, utilizing GraphPad Prism. Differences were considered significant at p-values ≤ 0.05.

