## Supplementary information for "A BRCT Domain-Containing Protein Induced in Early Phagocytosis Plays a Crucial Role in Mucorales Pathogenesis"

A

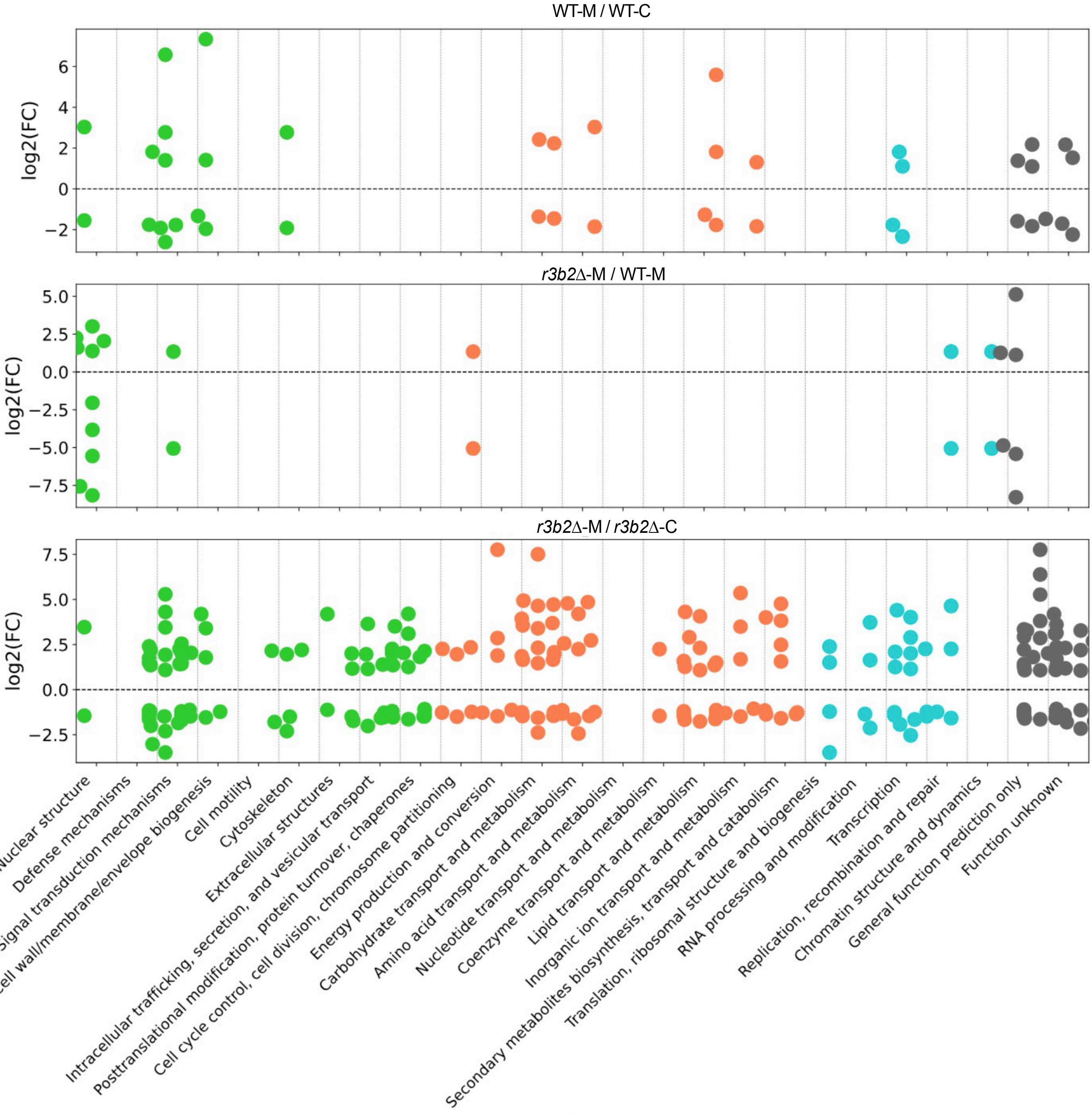
WT-M / WT-C
log2(FC)
r3b2Δ-M / WT-M
log2(FC)
r3b2Δ-M / r3b2Δ-C
log2(FC)
Nuclear structure
Defense mechanisms
Signal transduction mechanisms
Cell wall/membrane/envelope biogenesis
Cell motility
Cytoskeleton
Extracellular structures
Intracellular trafficking, secretion, and vesicular transport
Posttranslational modification, protein turnover, chaperones
Cell cycle control, cell division, chromosome partitioning
Energy production and conversion
Carbohydrate transport and metabolism
Amino acid transport and metabolism
Nucleotide transport and metabolism
Coenzyme transport and metabolism
Lipid transport and metabolism
Inorganic ion transport and metabolism
Secondary metabolites biosynthesis, transport and catabolism
Translation, ribosomal structure and biogenesis
RNA processing and modification
Transcription
Replication, recombination and repair
Chromatin structure and dynamics
General function prediction only
Function unknown

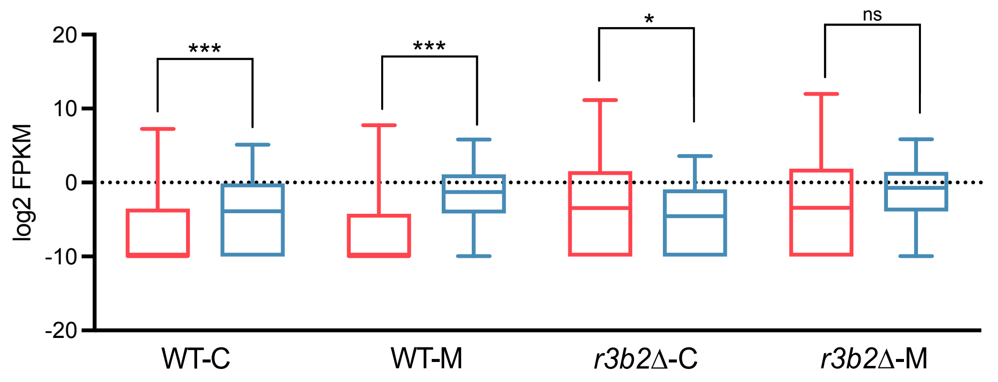
log2 FPKM
***
***
*
ns
WT-C
WT-M
r3b2Δ-C
r3b2Δ-M

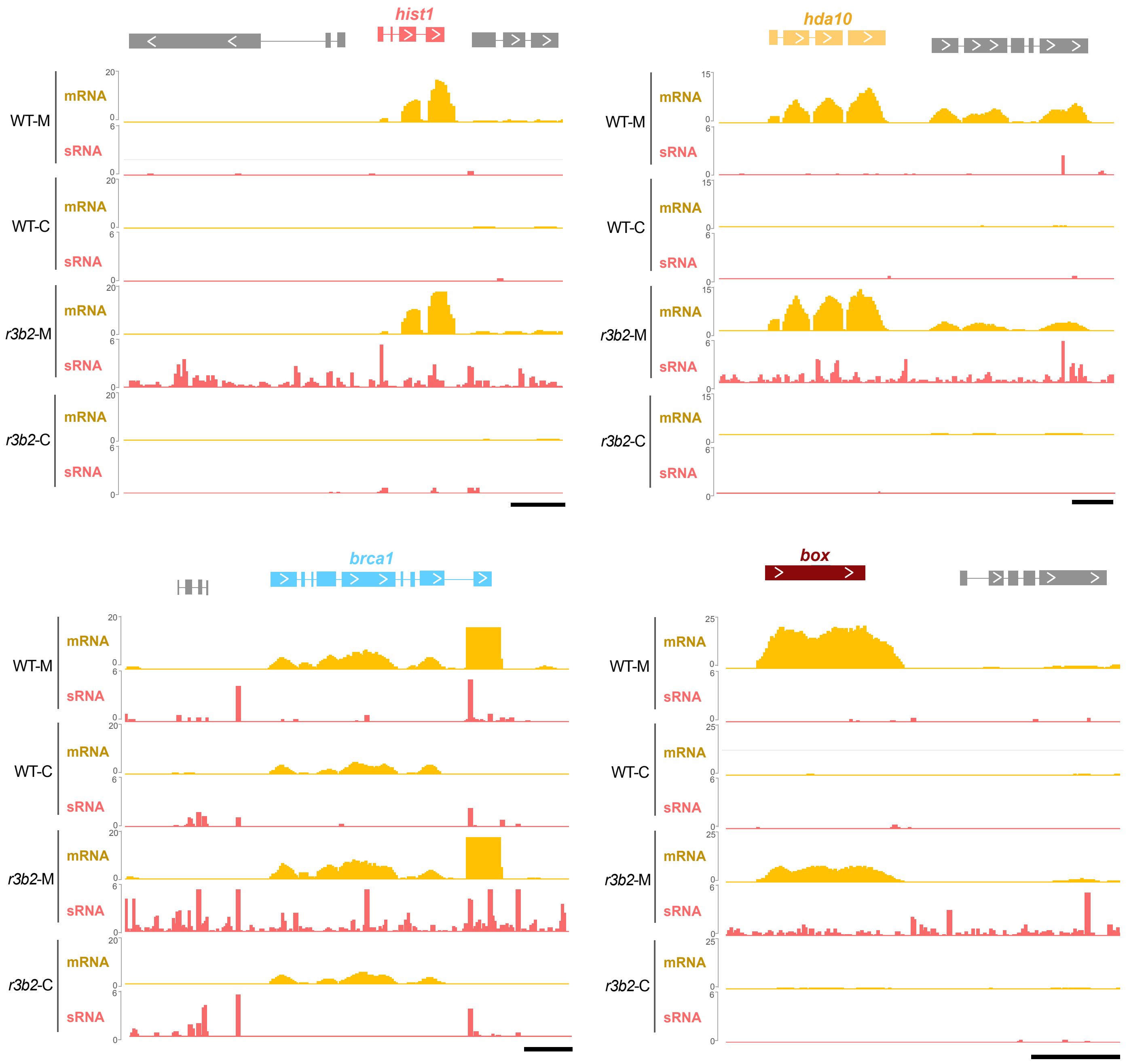

hist1
hda10
brca1
box
WT-M
WT-C
r3b2-M
r3b2-C
mRNA
sRNA

A

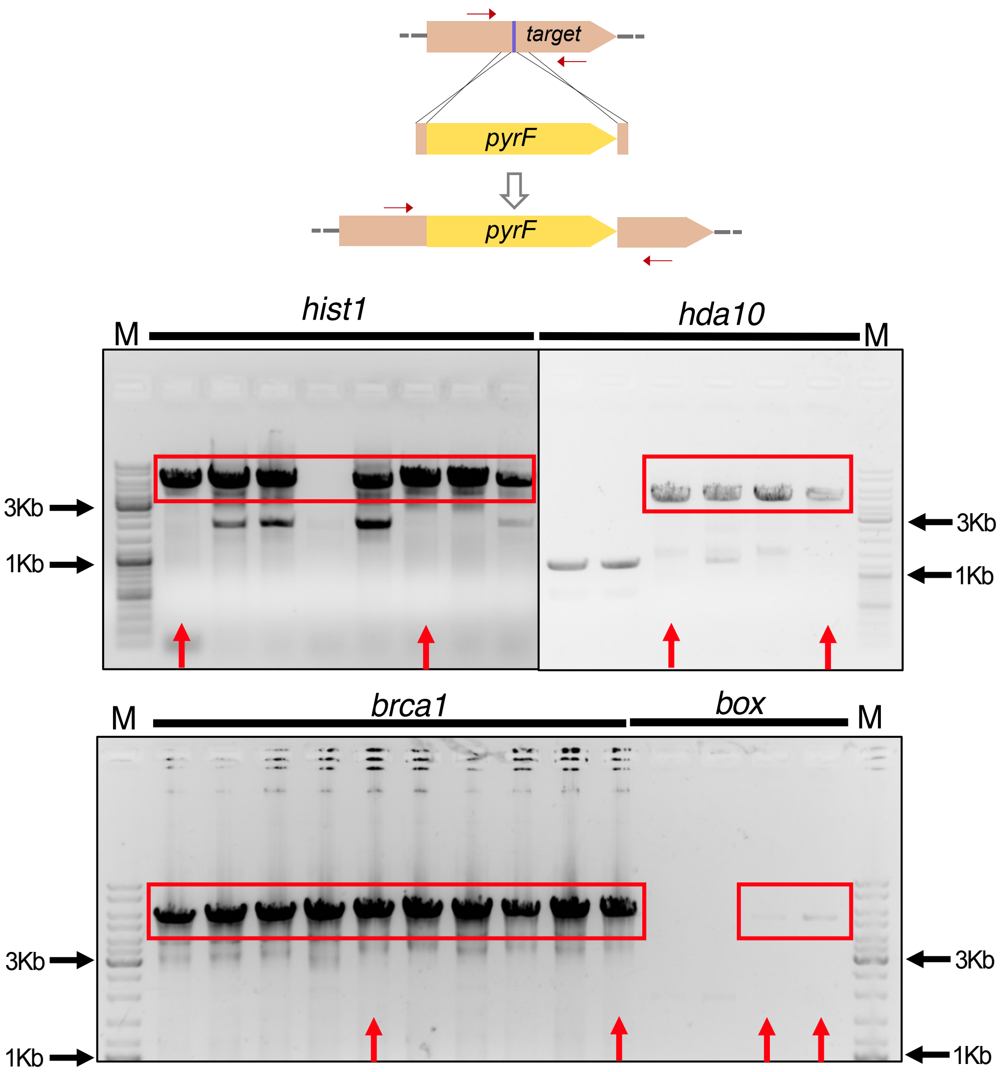
target
pyrF
pyrF
M
hist1
hda10
M
3Kb
1Kb
3Kb
1Kb
M
brca1
box
M
3Kb
1Kb
3Kb
1Kb

B

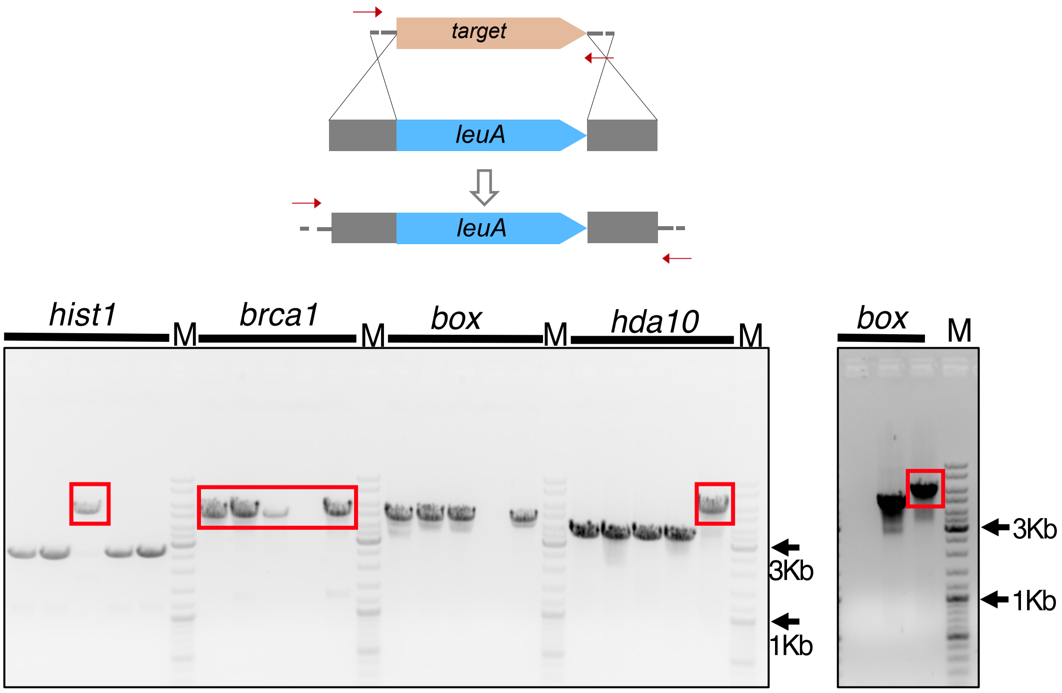
target
leuA
leuA
hist1
M
brca1
M
box
M
hda10
M
3Kb
1Kb
box
M
3Kb
1Kb

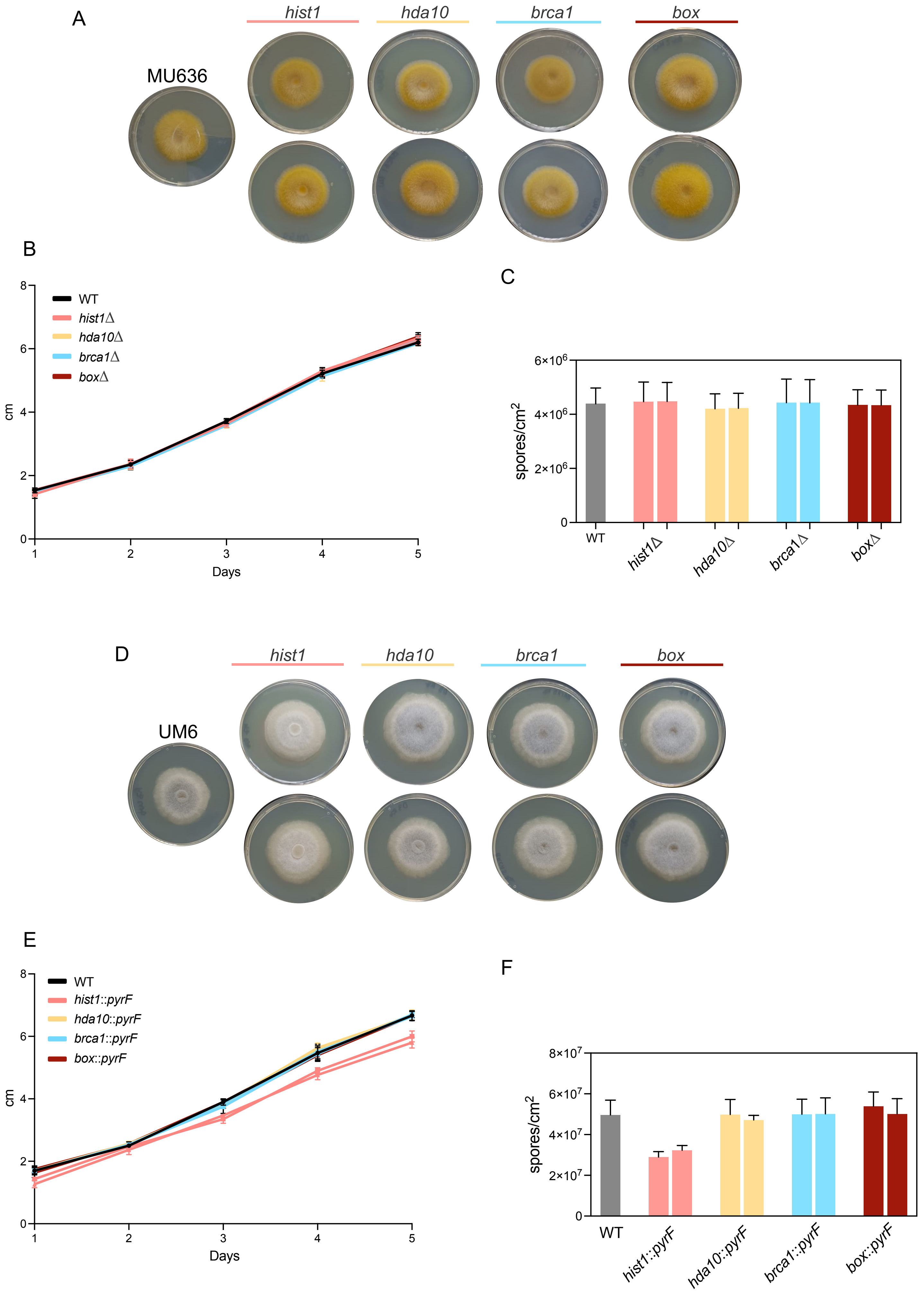
A
hist1
hda10
brca1
box
MU636
B
WT
hist1Δ
hda10Δ
brca1Δ
boxΔ
cm
Days
C
spores/cm2
6×10^6
4×10^6
2×10^6
0
WT
hist1Δ
hda10Δ
brca1Δ
boxΔ
D
hist1
hda10
brca1
box
UM6
E
WT
hist1::pyrF
hda10::pyrF
brca1::pyrF
box::pyrF
cm
Days
F
spores/cm2
8×10^7
6×10^7
4×10^7
2×10^7
0
WT
hist1::pyrF
hda10::pyrF
brca1::pyrF
box::pyrF

A

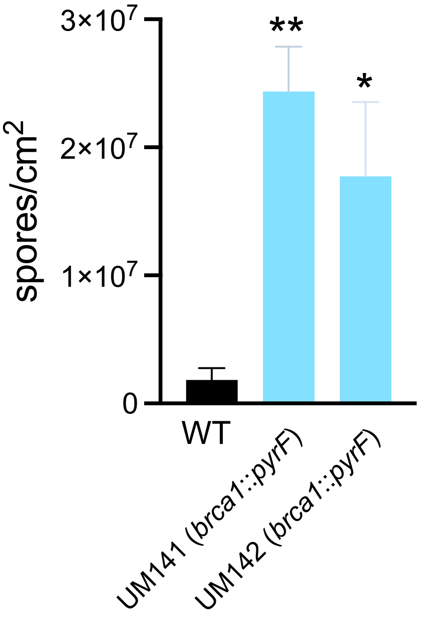

3×10^7
2×10^7
1×10^7
0
spores/cm^2
**
*
WT
UM141 (brca1::pyrF)
UM142 (brca1::pyrF)


B

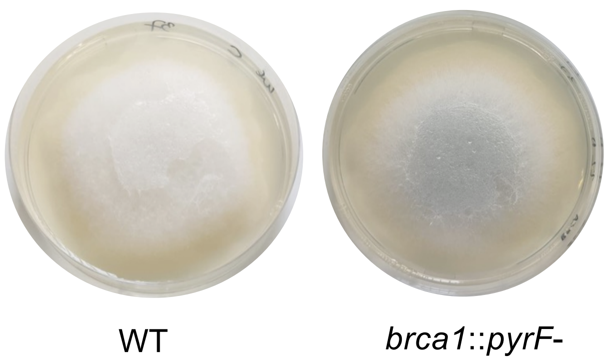

WT
brca1::pyrF-

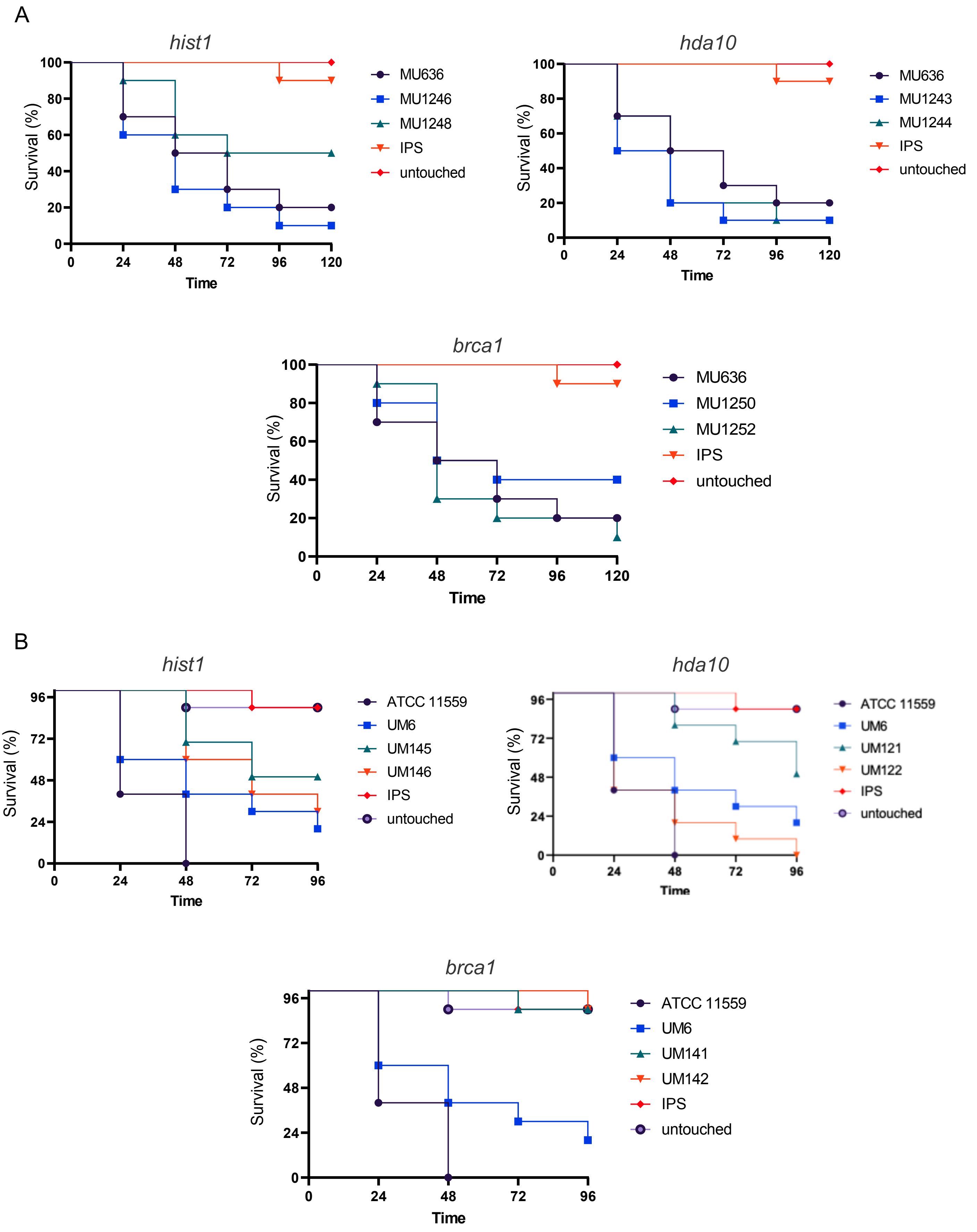
A
hist1
Survival (%)
100
80
60
40
20
0
0
24
48
72
96
120
Time
MU636
MU1246
MU1248
IPS
untouched
hda10
Survival (%)
Time
MU636
MU1243
MU1244
IPS
untouched
brca1
Survival (%)
Time
MU636
MU1250
MU1252
IPS
untouched
B
hist1
Survival (%)
96
72
48
24
0
Time
ATCC 11559
UM6
UM145
UM146
IPS
untouched
hda10
Survival (%)
Time
ATCC 11559
UM6
UM121
UM122
IPS
untouched
brca1
Survival (%)
Time
ATCC 11559
UM6
UM141
UM142
IPS
untouched

Table 1. Fungal strains.

| Strain | Genotype | Description | Organism | Source |
|---|---|---|---|---|
| ATCC11559 | WT | WT strain | *R. microsporus* | Lax et al. 2021 |
| UM33 | *pyrF-, leuA-* | Avirulent Control. Used as a receptor strain for the mutant generation | *R. microsporus* | Lax et at. 2025 |
| UM6 | *pyrF+, leuA-* | Virulent strain | *R. microsporus* | Lax et al. 2021 |
| UM121 | *leuA-, hda10::pyrF* | Mutant in which the *hda10* gene has been disrupted with the *pyrF* gene. Derived from UM33 | *R. microsporus* | This work |
| UM122 | *leuA-, hda10::pyrF* | Mutant in which the *hda10* gene has been disrupted with the *pyrF* gene. Derived from UM33 | *R. microsporus* | This work |
| UM141 | *leuA-, brca1::pyrF* | Mutant in which the *brca1* gene has been disrupted with the *pyrF* gene. Derived from UM33 | *R. microsporus* | This work |
| UM142 | *leuA-, brca1::pyrF* | Mutant in which the brca1 gene has been disrupted with the *pyrF* gene. Derived from UM33 | *R. microsporus* | This work |
| UM143 | *leuA-, box::pyrF* | Mutant in which the *box* gene has been disrupted with the pyrF gene. Derived from UM33 | *R. microsporus* | This work |
| UM144 | *leuA-, box::pyrF* | Mutant in which the *box* gene has been disrupted with the *pyrF* gene. Derived from UM33 | *R. microsporus* | This work |
| UM145 | *leuA-, hist1::pyrF* | Mutant in which the *hist1* gene has been disrupted with the *pyrF* gene. Derived from UM33 | *R. microsporus* | This work |
| UM146 | *leuA-, hist1::pyrF* | Mutant in which the *hist1* gene has been disrupted with the *pyrF* gene. Derived from UM33 | *R. microsporus* | This work |
| R7B | *pyrG+, leuA-* | WT virulent | *Mucor lusitanicus* | Pérez-Arques et al 2019 |
| NRRL3631 | *pyrG+, leuA+* | WT avirulent | *Mucor lusitanicus* | Pérez-Arques et al 2020 |
| MU412 | *pyrG+, leuA-* | Homokaryotic mutant | *Mucor lusitanicus* | Trieu et al. 2015 |
| MU636 | *pyrG+, leuA-* | WT virulent | *Mucor lusitanicus* | Pérez-Arques et al 2019 |
| CBS277.49 | *pyrG+, leuA+* | WT virulent (prototroph) | *Mucor lusitanicus* | Pérez-Arques et al 2020 |
| MU1243 | *pyrG+, leuA+, hda10Δ* | Homokaryotic mutant derived from MU636 in which the 168144 (*hda10*) gene has been deleted. Checked by PCR | *Mucor lusitanicus* | This work |
| MU1244 | *pyrG+, leuA+, hda10Δ* | Homokaryotic mutant derived from MU636, in which the 168144 (*hda10*) gene has been disrupted. Checked by PCR | *Mucor lusitanicus* | This work |
| MU1246 | *pyrG+, leuA+, hist1Δ* | Homokaryotic mutant derived from MU636 in which the 83400 gene (*hist1*) has been disrupted. Checked by PCR | *Mucor lusitanicus* | This work |

| | | | | |
|---|---|---|---|---|
| MU1248 | *pyrG+, leuA+, hist1Δ* | Homokaryotic mutant derived from MU636 in which the 83400 gene (*hist1*) has been disrupted. Checked by PCR | *Mucor lusitanicus* | This work |
| MU1250 | *pyrG+, leuA+, brca1Δ* | Homokaryotic mutant derived from MU636 in which the 113714 (*brca1*) gene has been disrupted. Checked by PCR | *Mucor lusitanicus* | This work |
| MU1252 | *pyrG+, leuA++, brca1Δ* | Homokaryotic mutant derived from MU636 in which the 113714 (*brca1*) gene has been disrupted. Checked by PCR | *Mucor lusitanicus* | This work |
| MU1254 | *pyrG+, leuA+, boxΔ* | Homokaryotic mutant derived from MU636 in which the 104613 (*box*) gene has been disrupted. Checked by PCR | *Mucor lusitanicus* | This work |
| MU1255 | *pyrG+, leuA+, boxΔ* | Homokaryotic mutant derived from MU636 in which the 104613 (*box*) gene has been disrupted. Checked by PCR | *Mucor lusitanicus* | This work |

*Table 2. Primers and gRNAs used in the mutant generation in Mucor lusitanicus and Rhizopus microsporus.*

| | Amplified gene | 5'3' |
|---|---|---|
| q-RT-PCR | 89889 *ltr-transposon*_Fw | GGACGCAATCACTGAACTGC |
| | 89889 *ltr-transposon*_Rv | TTGGCGGCGTAATTCCTTCAG |
| | 165343 *permease*_Fw | TTGAGAACCTGGCGGATGAG |
| | 165343 *permease*_Rv | GACCTTGGCCGATAACACCA |
| | 83143 *prd*_Fw | TGTCTCGCCTTGCTATCCTG |
| | 83143 *prd*_Rv | CAGTTGTCGCCGAGTCCATATC |
| | 113714 *brca1*_Fw | CATCATCGCAACAGCAGCAG |
| | 113714 *brca1*_Rv | TTCGGTTCTCACGTCCAGAC |
| | 104613 *boxC*_Fw | TCAGCCAGTACGACACATTC |
| | 104613 *boxC*_Rv | GGTGATGAAGCGGTTGATGG |
| | 83400 *hist1*_Fw | CTCGTCCTGCCATCAAGAAG |
| | 83400 *hist1*_Rv | CACGTCGAATGGCACCAGAG |
| | 168144 *hda1*_Fw | CGCTTAACAGAGGCCATTGC |
| | 168144 *hda1*_Rv | GAGCAACAACAGATGCATTG |
| | BRCA1_1Kb_UP_Fw | CTTGAAGGCAAGATCCTCCTG |
| | BRCA1_1Kb_UP_Rv | CtGAATAGAGTTGGTAGGGAGCATACGTACTCGGATAT AGCTTGGAATAATAAG |
| | BRCA1_1Kb_Down_Fw | CattttGTACGATTCTGGTCAACTCGACCTTAAATCTCCA CATCTAAATGAAC |
| | BRCA1_1Kb_Down_Rv | GATTGTGACAAAAACCGCATG |
| | boxC_1Kb_UP_Fw | GCCCAGTAATTGGCATTACATG |
| | boxCHMG_1Kb_UP_Rv | ctGAATAGAGTTGGTAGGGAGCATACGTACTCGTCATG ATCATTTATATCTGATTTAG |
| | boxC_1kb_Down_Fw | CattttGTACGATTCTGGTCAACTCGACCTTGAAGTCAAC TCTTGGGTGATTTTG |
| | boxC_1kb_Down_Rv | TAAAGCGTGCTCCTGTAGTC |
| | hist1_1Kb_UP_Fw | TACGAGTGGCTGCTTGAGTAGAG |
| | hist1_1Kb_UP_Rv | CtGAATAGAGTTGGTAGGGAGCATACGTACTCGTGAG TCGATAATATGAAGAATG |

| | | |
|---|---|---|
| Phagocytosis-related mutants in *Mucor lusitanicus* | hist1_1Kb_Down_Fw | cattttGTACGATTCTGGTCAACTCGACCTCCACTGTCAAAGCTTAATGC |
| | hist1_1Kb_Down_Rv | TTGTTCAGCTTGGGTAGGTC |
| | hda1_1Kb_UP_Fw | CTTGCATTGTTTGGAATCTTC |
| | hda1_1Kb_UP_Rv | ctGAATAGAGTTGGTAGGGAGCATACGTACTCGTTTGGTGTTGCCGAGTTGAAG |
| | hda1_1Kb_Down_Fw | cattttGTACGATTCTGGTCAACTCGACCTTGAACAGCAATGAATAAAAAGC |
| | hda1_1Kb_Down_Rv | ACAATCCAGGCACAGTGACAAG |
| | brca1_locusM_Fw | ACTTCTTGGCCTCTTCCATC |
| | brca1_locusM_Rv | GTAATAAGGCAACCTTAAGCAC |
| | boxC_locusM_Fw | GTCTGACCGTGTGCACAGTC |
| | boxC_locusM_Rv | GAATGGTTTCAGCATCGTATTC |
| | hist1_locusM_Fw | TGCAGCAAAGAGTCCCTTCTAC |
| | hist1_locusM_Rv | TTGGTATTTGATTTTGTTCAG |
| | hda1_locusM_Fw | GCATCTATGCGGTTGTTTGAG |
| | hda1_locusM_Rv | GATCTTCAACAAGATGGGCAAC |
| | leuAW1_F | CGCCTCATTGAGTCACTGCCAG |
| | leuAW3_R | GGAACAACCAGCCTCTCTCC |
| | LeuAFow3kb-Eco10I-XhoI | CCCCCTCGAGTACGTATGCTCCCTACCAACTCTATTC |
| | LeuARev3Kb-PstI | CCCCCTGCAGGTCGAGTTGACCAGAATCGTAC |
| Phagocytosis-related mutants in *Rhizopus microporus* | pyrF_F_hist1_Fw | GGTGCCCACTTTGATACCCAAGTTGCCGCTGCTATCAATCCTCCATAAGAATTTGACAG |
| | pyrF_R_hist1_Rv | CTTTAGGAAGTTCAAAGATGCCCTTGGCATGGCCACGCTGATAAAACGAAGATGTGGCTGTC |
| | pyrF_F_hda1_Fw | TCATTCAAGCCAATGACTATTCCTTAAAACCAATCCTTTCCTCCATAAGAATTTGACAG |
| | pyrF_R_hda1_Rv | TCTGTTAAAAACTGAATATAGTCTGATGGATGAACCTCTGATAAAACGAAGATGTGGCTGTC |
| | pyrF_F_boxC_Fw | GAGCTGGTTAAAGGATGGAGTGAATGGTGGCCCTTCATTCCTCCATAAGAATTTGACAG |
| | pyrF_R_boxC_Rv | TGACTTTTTTGATAACCAGTTGACCAACACATCCATAGTGATAAAACGAAGATGTGGCTGTC |
| | pyrF_F_brca1_Fw | CCAGAAAACAAAATATTAGAAGGTGTAGTTGCCTGTTTTCCTCCATAAGAATTTGACAG |
| | pyrF_R_brca1_Rv | AACGTGATCAAGTACATATTGTTACTACCTAACATCTATGATAAAACGAAGATGTGGCTGTC |
| | hist1_Rz_locus_Fw | TTGATCAGAATGACCTTGGC |
| | hist1_Rz_locus_Rv | ATCCTATGGTAAGGCCTTCTC |
| | hda1_Rz_locus_Fw | ATTAAGTATCTGTAATGAAG |
| | hda1_Rz_locus_Rv | TCCATAAAGCCCTTTAATAC |
| | brca1__Rz_locus_Fw | GTTACATGAAGGCTATCAAG |
| | brca1__Rz_locus_Rv | CACAACGGATGCGGATGATG |
| | boxC_Rz_locus_F | CAGTGATAAAAGGCGCGTAG |
| | boxC_Rz_locus_R | TTATAGTTTATTCAATTGCTC |

*Table 1. gRNAs used in the mutant generation in Rhizopus microsporus.*

| | Target gene | 5'3' |
|---|---|---|
| gRNAs for the phagocytosis related genes in *Rhizopus microsporus* | gRNA_hist1 | GTTGCCGCTGCTATCAAGCG |
| | gRNA_hda1 | TTCCTTAAAACCAATCCTTG |
| | gRNA_boxC | GAATGGTGGCCCTTCATCTA |
| | gRNA_brca1 | TTACTACCTAACATCTAAAC |
